# Polygenic architecture of brain structure and function, behaviors, and psychopathologies in children

**DOI:** 10.1101/2024.05.22.595444

**Authors:** Yoonjung Yoonie Joo, Eunji Lee, Bo-Gyeom Kim, Gakyung Kim, Jungwoo Seo, Jiook Cha

## Abstract

The human brain undergoes structural and functional changes during childhood, a critical period in cognitive and behavioral development. Understanding the genetic architecture of the brain development in children can offer valuable insights into the development of the brain, cognition, and behaviors. Here, we integrated brain imaging-genetic-phenotype data from over 8,600 preadolescent children of diverse ethnic backgrounds using multivariate statistical techniques. We found a low-to-moderate level of SNP-based heritability in most IDPs, which is lower compared to the adult brain. Using sparse generalized canonical correlation analysis (SGCCA), we identified several covariation patterns among genome-wide polygenic scores (GPSs) of 29 traits, 7 different modalities of brain imaging-derived phenotypes (IDPs), and 266 cognitive and psychological phenotype data. In structural MRI, significant positive associations were observed between total grey matter volume, left ventral diencephalon volume, surface area of right accumbens and the GPSs of cognition-related traits. Conversely, negative associations were found with the GPSs of ADHD, depression and neuroticism. Additionally, we identified a significant positive association between educational attainment GPS and regional brain activation during the N-back task. The BMI GPS showed a positive association with fractional anisotropy (FA) of connectivity between the cerebellum cortex and amygdala in diffusion MRI, while the GPSs for educational attainment and cannabis use were negatively associated with the same IDPs. Our GPS-based prediction models revealed substantial genetic contributions to cognitive variability, while the genetic basis for many mental and behavioral phenotypes remained elusive. This study delivers a comprehensive map of the relationships between genetic profiles, neuroanatomical diversity, and the spectrum of cognitive and behavioral traits in preadolescence.

## Introduction

Preadolescent childhood is a critical developmental period with a cascade of neurobiological changes. During this period, white matter volume increases substantially, and gray matter undergoes region-specific growth, peaking in the frontal and parietal lobes^1,2^. This dynamic interplay between genetics and environment not only shapes the maturing brain but also sets the trajectory for cognitive abilities and mental health outcomes in later life.

Cognitive functions and psychiatric conditions share common genetic and neurological roots^3,4^. The heritability of cognitive and psychological outcomes generally increases along the lifespan. For instance, genetics accounts for less than 25% of the variability in cognition in infancy, but up to 70% in adolescence^5^. Similarly, the heritability of various psychopathologies, including externalizing behaviors, depression, and anxiety, also appears to rise with age^6^. This evolving genetic influence may be attributed to developmental changes in gene expression or neuroanatomical alterations that render the brain more susceptible or resistant to genetic and environmental inputs at different life stages^7^. Given these substantial genetic influences, elucidating the genetic basis of neural and cognitive development is imperative to advance our understanding of the neurobiological foundations of cognition and psychopathology.

The joint analysis of multimodal genetic and brain imaging data could offer novel insights into interpreting the relationships between genetic variation and brain imaging-derived phenotypes (IDPs), and their relationship to cognitive and behavioral processes. Fueled by recent technological advances in high throughput data generation, computational approaches linking genetics to neuroimaging have become more central to the study of biological systems. Identifying which brain structures or functions are correlated with polygenic signals may allow us to better understand the biological relevance of their intricate intertwining, responsible for cognitive and psychological outcome development. However, existing studies do not sufficiently take account of the developmental stage of preadolescence^8^. While prior works have demonstrated significant connections between the brain and genomic components^9,10^, no studies have yet examined the polygenic influences on specific brain IDPs and behavioral outcomes during preadolescence.

We pivot from traditional analyses to employ sparse generalized canonical correlation analysis (SGCCA)^11,12^, an advanced multivariate technique optimized for high-dimensional data characteristic of neuroimaging and genetic datasets. SGCCA allows us to distill the essence of complex datasets, reducing feature space while preserving interpretability, a significant leap from earlier applications of canonical correlation analysis that lacked regional specificity in brain imaging-derived phenotypes (IDPs)^8,9,10^.

In a pioneering approach, we utilize a comprehensive suite of polygenic scores (GPS) to dissect the functional associations of genetic variations on brain IDPs and behavioral phenotypes, which could provide more interpretable and informative insights into trait variability. Our methodology contrasts with prior studies that relied on a narrow spectrum of GPS and could not capture the full genetic architecture influencing trait development^9^.

With the novel application of this multivariate method to the genetic and brain imaging data, we aimed to reveal the interconnected variations of genetic signals, the brain, and cognitive and psychological outcomes in the preadolescent brain. Our estimates of genetic heritability for multimodal brain features set the stage for SGCCA, enabling us to explore the covariation between IDPs and polygenic scores across 29 traits, potentially transforming our comprehension of the genetic orchestration of brain and behavioral development.

## Methods

### Study Participants

We used multimodal neuroimaging, DNA genotype data, and cognitive and behavioral metrics of 11,875 multiethnic children from the Adolescent Brain and Cognitive Development (ABCD) study. The ABCD study is a nationwide longitudinal cohort study to investigate the normative brain and cognitive development of preadolescent youths from ages 9 to 10 from 21 sites across the United States. The study provides a rich repository of multi-modal biomedical datasets such as DNA genotype, multimodal neuroimaging data, and matched cognitive, behavioral and clinical data. The ethical approval for the study was obtained from Seoul National University Institutional Review Board (IRB).

### Genetic Data

The ABCD study collected and genotyped the participants’ saliva samples at Rutgers University Cell and DNA Repository (RUCDR) using Affymetrix SmokeScreen Array consisting of 733,293 single nucleotide polymorphisms (SNPs). We applied standard PLINK filters for genotype call rate (<95% removed), sample call rate (<95% removed), and minor allele frequency (MAF <1% removed). Genotypes were imputed toward 1000 Genome phase3 reference panel^10^ using the Michigan Imputation Server^11^, and the output was phased with Eagle v2.4^12^ (a total of 12,046,090 SNPs). After imputation, we additionally removed any inferior SNPs with low imputation score (INFO score <0.4), genotype call rate <95%, Hardy-Weinberg Equilibrium p-value <1E-20 for diverse ethnical population, sample missingness >5%, minor allele frequency (MAF) <0.5%, and extreme heterozygosity (over three standard deviations of the population mean). After this step, 11,221,810 variants remained.

### Relatedness Inference

Since the ABCD study participants have diverse ethnic backgrounds and genetic ancestries, we performed an additional QC process to thoroughly address potential population stratification due to genetic relatedness and ancestry admixture. To remove potential confounds of the family structure, we identified genetically unrelated individuals and computed the ancestrally informative principal components (PCs) of their genotype data using *SNPRelate* R package^13^. Two rounds of principal component analysis (PCA) were conducted using the PC-Air algorithm that is robust to familial or cryptic relatedness. We first obtained initial estimates of pairwise kinship coefficients using KING-robust algorithm using a pruned set of independent genetic variants using LD threshold of *r*^2^ < 0.1. We identified genetically related individuals with closer than 3^rd^ degree relatives (kinship threshold = 2^(-9/2)^, which defines anyone less than first cousins as ‘unrelated’) and only retained one individual per related pair and kept the excluded individual as an independent validation set (n=1,814). PC-Air identified the unrelated set of 8,845 individuals based on the pairwise kinship estimates, and systematically computed PC of their genotype data. PC-Relate was used to compute new kinship estimates adjusting for ancestry, which is robust to population structure and admixture in estimating genetic relatedness. The 2^nd^ round of PC-Air computed accurate PCs with the modified unrelated set. We further excluded 88 participants whose projected PCs had the Mahalanobis distances greater than 6SDs from further analysis. In conclusion, the final set of 8,620 unrelated individuals was used for the main analysis. The excluded 1,814 individuals from the relatedness analysis were additionally assessed for their 3^rd^ degree relatedness and the final unrelated set of 1,579 individuals were kept for the independent validation set for GPS optimization and CCA hyperparameter tuning.

### Construction of Genome-Wide Polygenic Scores (GPSs)

To estimate polygenic liability of complex traits for each individual, we constructed 29 different cognitive, psychological, and psychiatric traits: attention-deficit/hyperactivity disorder (ADHD)^14^, cognitive performance (CP)^15^, educational attainment (EA)^15^, major depressive disorder (MDD)^16^, insomnia^17^, snoring^17^, intelligence quotient (IQ)^18^, post-traumatic stress disorder (PTSD)^19^, depression^20,21^, body mass index (BMI)^22,23^, alcohol dependence (alcohol use)^24^, autism spectrum disorder (ASD)^25^, automobile speeding propensity (ASP)^26^, bipolar disorder^27^, cannabis during lifetime (cannabis use)^28^, ever smoker (smoking status)^26^, shared effects on five major psychiatric disorders (cross disorder)^29^, alcoholic consumption per week (drinking)^26^, eating disorder^30^, neuroticism^31^, obsessive-compulsive disorder (OCD)^32^, first principal components of four risky behaviors (PC of risky behaviors)^26^, general risk tolerance^26^, schizophrenia^33,34^, worrying^31^, subjective wellbeing^35^, general happiness, and general happiness for health (happiness-health) and meaningful life (happiness-life). We collected the GWAS summary statistics of 29 complex traits from publicly available resources listed in **Table S1.** For better cross-population polygenic prediction, PRS-CSx^36^, a recently developed Bayesian polygenic modeling technique, was used to construct the GPSs for our multi-ancestry study participants. The method is known for its effective posterior inference algorithm considering population-specific allele frequencies and LD patterns by adopting a shared continuous shrinkage prior. The LD reference panel of European-ancestry (EUR) and Admixed Americans (AMR) from the 1000 Genome Project phase 3 was utilized in accordance with the discovery GWAS sample for GPS construction. Though PRS-CSx could automatically estimate the parameter, we applied small-scale grid search of global shrinkage parameter for whose target phenotype is available within the study dataset for better predictive performance.

### Validation of Genome-Wide Polygenic Scores (GPSs)

To estimate the optimal GPS, we manually tuned the best-performing global shrinkage hyperparameter to maximize the explained variance of the GPSs on the corresponding phenotype available in the ABCD datasets. For example, ADHD behavioral measure of the ABCD study was used to choose the best version of the GPS of ADHD with optimal hyperparameter. In this way, we optimized GPSs of 14 traits by choosing the optimal global shrinkage hyperparameter (φ, phi) in an independent held-out validation set of 1,579 unrelated participants, which were removed during the QC process of relatedness analysis. We examined the effect size and significance of the GPS variable in linear regression model with different phi values results from the small-scale grid search (φ = 1e-6, 1e-4, 1e-2, 1). Each target outcome variable was regressed on the relevant GPS, sex, top 10 principal components (PCs) of genotype data, and genetic ancestry. We chose the final φ value of the regression model maximizing effect size (β coefficient) of the GPS and *R^2^* of the model. For example, when the φ value is 1, the regression model would be:

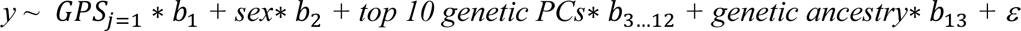

For five complex traits (i.e., BMI, PTSD, depression, schizophrenia, and alcohol dependence), whose GWAS summary statistics were available in both European- and non-European-ancestry participants, we built multiethnic GPSs combining the GWAS summary statistics of two or more ancestries by learning the optimal linear combination of the ancestry-specific GPSs that were used as predictors. For example, when the φ value of European-ancestry-based GPS is 1, and that of Admixed-American-based GPS is 1e-2, the regression model would be:

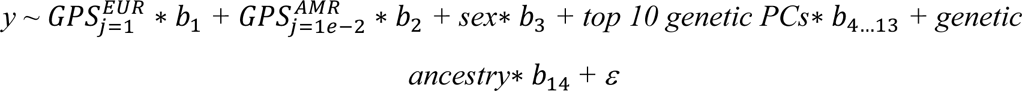

After optimal GPSs were built, we assessed the validity of final GPSs in 8,620 participants, the sample for our main analysis, adjusting for sex, age, and study site. Genetic ancestry was added as an additional covariate in tuning the multiethnic GPSs. For the 15 GWAS traits that do not have target outcome variables in the ABCD study, we performed pseudo-validation using PRS-CS-auto, in which the hyperparameter φ is automatically selected from data with a fully Bayesian approach^37^.

### Neuroimaging Data

We used different modalities of neuroimaging data available from the ABCD study, including structural magnetic resonance imaging (sMRI), diffusion connectivity measures (dMRI), resting-state functional MRI (rs-fMRI), and task functional MRI data (task fMRI). We processed T1-weighted (T1w) 3D structural MRI using FreeSurfer v6.0 (https://surfer.nmr.mgh.harvard.edu). After excluding global brain measures (i.e., total intracranial volume (TIV)), we used cortical and subcortical surface area, volume, thickness, and mean curvature in the analysis. In addition, we applied MRtrix3^38^ for whole-brain white matter tract estimation and individualized connectome generation for diffusion spectrum imaging. For connectivity matrices, we used fractional anisotropy (FA) and streamline counts related to fiber connection strength. Computation for diffusion MRI was done using the supercomputers at Argonne Leadership Computing Facility Theta and Texas Advanced Computing Center Stampede2. For the resting-state functional MRI acquisition, we used pair-wise correlation coefficients between each cortical and subcortical region of interest (ROI) and network. For task fMRI, we used three tasks: the stop-signal task (SST)^39^, the emotional version of N-back (N-back)^40^, and the monetary incentive delay task (MID)^41^. To ensure data quality, we excluded individuals who had more than 10% missing data in any of the neuroimaging modalities, resulting in the removal of 559 children from the analysis. Additionally, we removed low-frequency variables that exhibited zero variance or had less than 100 observations among the participants.

### Non-imaging Measures and Data Processing

In addition to investigating the associations between brain IDPs and polygenic scores, we examined the baseline non-imaging measures of mental and physical health, neurocognition, culture and environment measures of the ABCD participants to explore their links with polygenic liability^42^. For assessing the mental wellbeing of the children, we used a parent version of Kiddie-Structured Assessment for Affective Disorders and Schizophrenia for DSM-5 (KSADS-5) and Child Behavioral Checklist (CBCL), which assess a wide range of emotional and behavioral aspects of the youths in the previous six months, including their mood, psychosis, anxiety, suicidality, behavioral and sleep problems. We also used Parent General Behavioral Inventory – mania (PGBI), Prodromal Psychosis Scale (PPS), Behavioral Inhibition/behavioral Approach System (BIS/BAS) scales, and UPPS-P for children (UPPS). For assessing physical well-being, we retrieved a lifetime medical history, head injury experiences, developmental medical records of the youth, experiences about sleep problems, pain, and exercises. For assessing neurocognition, we used the NIH Toolbox measurement, which consists of seven domain-specific tasks assessing episodic memory, executive function, attention, working memory, processing speed, and language abilities of the children^43,44^. We used age-uncorrected task scores of the NIH Toolbox and included age as covariate for further analysis. Also, we considered school attributes, population density, neighborhood walkability, and area deprivation indices of the children’s homes and neighborhood as part of the culture and environment phenotype block, which were reported directly by the youth or parent/caregiver. The description of all phenotypic variables included in our analyses is in **Table S2**.

Except for a few mental health phenotypes, the average missingness of the non-imaging phenotype block was around 3% and it was manually imputed. We imputed the missing values with the mode for categorical variables and the median for continuous variables (**Table S3**). We also left bothering scales of prodromal psychosis not imputed, coded only when one has at least one prodromal psychotic symptom. We identified and removed any non-imaging variables with high correlation with other variables (Pearson’s r > 0.95) or zero variance, resulting in the removal of 6 variables related to mental wellbeing and 1 cultural variable.

### SNP-Based Heritability

We estimated the SNP-based heritability of imaging and non-imaging phenotypes and the genetic correlation among these traits in children. The genetic relationship matrix (GRM) was generated from imputed and autosomal SNPs using GCTA v1.93, quantifying addictive genetic relatedness between pairs of participants. GCTA’s Restricted Maximum Likelihood algorithm used the GRM to estimate the variance explained by all SNPs for each trait. The significance of the estimates was determined using a likelihood ratio test, which compared the likelihood of the alternative to that of the null hypothesis.

During the preprocessing step, we identified and excluded features with zero variance and outliers greater than 5 median absolute deviations (MAD) from the median. All the phenotype data were quantile transformed.

In testing heritability, we included age, sex, age², age*sex, age²*sex, self-reported ethnicity, genetic-based ethnicity, the first ten principal components (PCs) of genotype data, and data collection sites as covariates for each phenotype^8^. We also applied additional imaging modality-specific covariates for the different modality of brain IDPs: 1) For structural MRI, signal-to-noise ratio of T1 and T2 brain mask across all OK scans, 2) For diffusion MRI, signal-to-noise ratio of b=0 image for all OK scans, mean intensity within brain mask averaged across all OK scans, the total number of censored slices in all frames for all OK scans, 3) For resting-state fMRI, signal to noise ratio within the brain of all OK scans, mean framewise displacement in mm, 4) For task-based fMRI (SST, N-back, MID): signal to noise ratio within the brain of all OK scans, average framewise displacement in mm. To validate the reliability of our results, we performed the analysis in the participants with European and multiethnic ancestries, respectively.

### Sparse Generalized Canonical Correlation Analysis (SGCCA)

SGCCA is a useful approach to explore the multivariate associations between high dimensional datasets and to discover subsets of canonical variables from each block that are significantly influential in their correlation with the other block. In SGCCA, the sets of variables observed on the same individuals were defined as blocks. We conducted SGCCA^45^ between the genomic block comprising the GPSs of 29 traits on the one hand, and each of seven neuroimaging data modality blocks (sMRI, dMRI (FA, streamline count), resting-state fMRI, and task fMRI (MID, SST, N-back)) and a non-imaging phenotype block on the other hand. To adjust for potential confounding effects, we took the residuals of the outcome variables after regressing out the baseline age, sex, age^2^, age*sex, age^2^*sex, self-reported ethnicity and data collection site. We performed the analysis using the *RGCCA* R package *(Regularized Generalized Canonical Correlation Analysis)*^46^. First, the level of regularization (sparsity) was determined using permutation testing (100 permutation) by randomly shuffling the participants from the held-out validation set. After tuning the optimal regularization for each modality, the main SGCCA analysis was performed in the main study participants. We built five components for each block. The number of components was chosen after preliminary experiments with different number of canonical components, which maximizes the cumulative variance explained of the overall imaging and non-imaging modalities. To assess the statistical significance of each component, we conducted a permutation test with 1,000 permuted datasets. The p-values of each component were estimated based on the number of permuted datasets having covariance greater than the covariance from the original dataset. All p-values were adjusted with FDR correction. Covariance and Pearson correlation coefficients between blocks were presented in the result section.

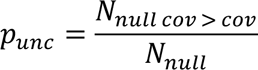

We calculated the loadings of the GPS variables and corresponding variables (e.g., phenotypes). To examine the reliability of loadings estimated from SGCCA, the bootstrap approach was used (bootstrap n=1,000) to evaluate the mean, variance, confidence intervals of the estimates across the bootstrap samples. To generate stable results, we averaged the canonical weights across ten folds. The statistical significance (p-value) of each loading was estimated based on the following null hypothesis: Fisher transformation of loading will follow normal distribution of *N*(0, *σ*^2^), where *σ* is estimated standard deviation of Fisher transformed loading from bootstrap samples. To assess the cross-population generalizability of our findings, we repeated the analysis in the entire participants with multiple genetic ancestries that mostly includes individuals of European-ancestry, but also the subjects of Asian-ancestry (n=112, 1.34%) and African-ancestry (n=1,361, 16.3%). Genetic ancestry was additionally included as covariates for multiethnic analysis.

### GPS-based Prediction of Phenotypes using Machine Learning Techniques

We examined the prognostic utility of the GPSs on cognitive and mental health outcomes in 6,555 European-ancestry and 8,620 multi-ancestry participants using several machine learning techniques. We employed the driverless artificial intelligence (AI) of H2O (version 1.10.1.2)^47^, which automates feature engineering, model ensemble and selection, optimization, and model interpretation to produce optimal machine learning-based prediction models. We manually chose 20 cognitive and psychological outcomes for our machine learning-based prediction models based on the results from the preceding CCA analysis, including six variables of NIH Toolbox neurocognitive measures (i.e., crystallized intelligence, oral reading recognition test, picture/vocabulary test, list sorting working memory test, total composite score, and fluid composite score), five behavioral problems (i.e., CBCL attention, rule-breaking, internalizing, externalizing, and total problems), and nine psychiatric conditions (i.e., KSADS-5 ADHD, parent and child reports of any psychiatric disorders, any depressive disorders, any anxiety disorders, and suicidal behaviors).

We trained several statistical and machine learning algorithms for prediction, including the constant model, generalized linear model (GLM), decision tree model, light gradient boosting machine (LightGBM), and extreme gradient boosting (XGBoost) with 5-fold cross-validation (CV). The automated feature engineering, early stopping, and automatic model tuning options of H2O were applied. We evaluated the model performance of regression analysis with root-mean-squared error (RMSE), mean absolute error (MAE), and *R^2^* (the squared Pearson correlation coefficient). For binary classification, accuracies were measured using precision, recall, and area under the receiver operating characteristic curve (AUC). Basic demographic variables (i.e., sex, age, household income, parental education, marital status, and research site) were used as input features of the baseline model in European-ancestry analysis, and we included genetic ancestry information as an additional covariate for multiethnic prediction. We evaluated the predictive performance of GPS-based model in comparison to the baseline model fitted with demographic variables. The relative importance of the input features in each model was reported by the absolute local Shapley values^48^ which indicates the degree to which each predictor contributes to the outcome.

## Results

### Study Participants

We initially considered multimodal data of 11,875 multiethnic participants from the ABCD study. After quality control procedures for brain imaging and genetic data, 6,555 European-ancestry participants were used for the primary analysis, and 8,620 multiethnic participants were used for complementary analysis. Descriptive statistics for each ancestry are presented in **Table 1**, **Figure S1,** and **Table S4**. We compared the estimated GPSs between European-ancestry and non-European-ancestry individuals, and all the GPSs showed no significant difference between different ancestries except for the IQ GPS (*P* = 0.0018) (**Table S4**).

### SNP-Based Heritability

We quantified the amount of variances in brain IDPs and psychological measures explained by all the imputed and QCed autosomal SNPs. Out of the 7,963 IDPs tested, 1,237 IDPs showed significant SNP heritability in 6,555 individuals of European ancestry (**Figure 1**; **Table 2**), ranging from 0.19 to 0.27. The average heritability of the brain IDPs (0.23) was higher than that of non-brain phenotypic measures (0.19).

**Figure 1.**
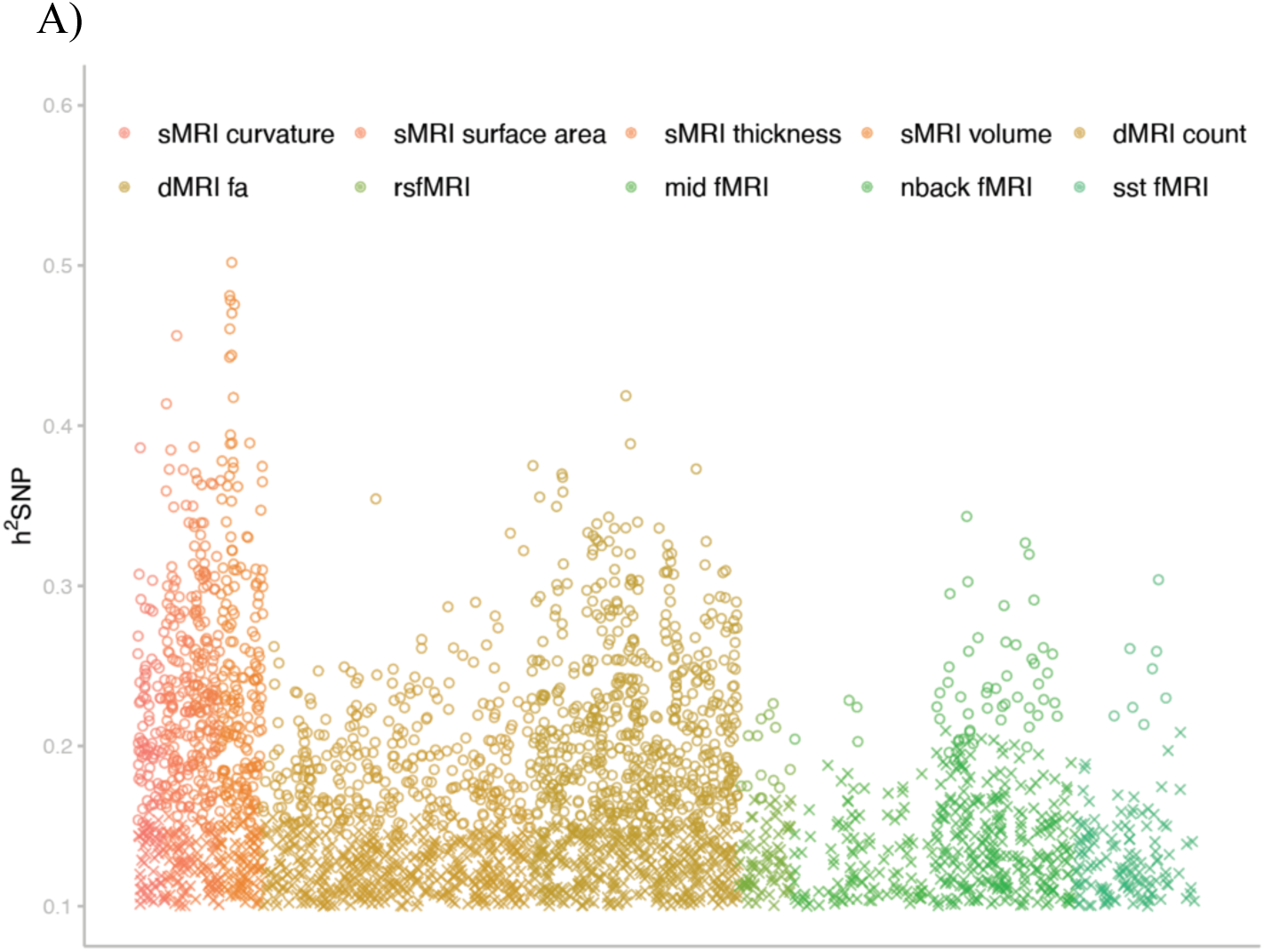
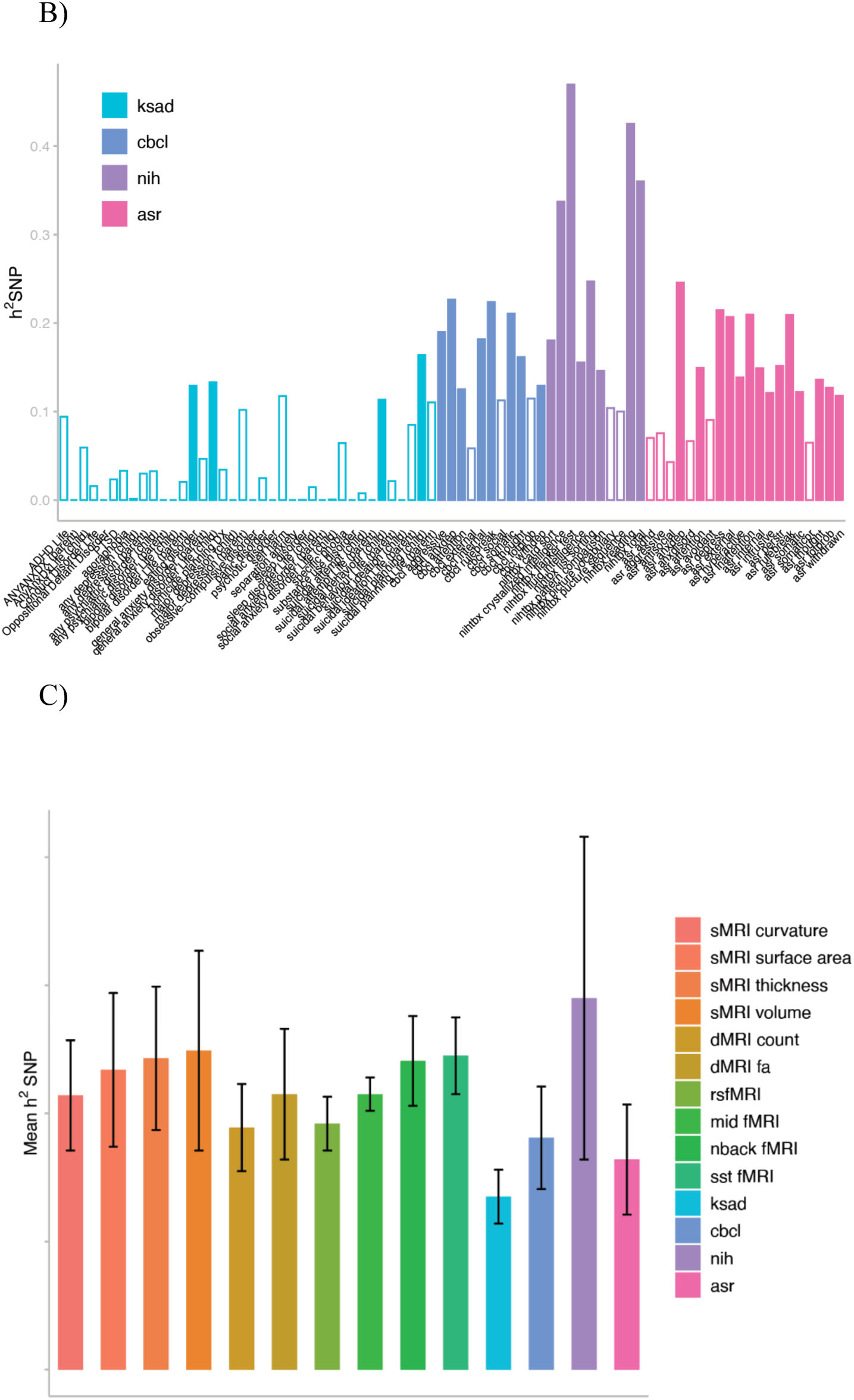
SNP heritability of brain IDPs and behavior phenotypes in European ancestry. A, Estimated SNP heritability of brain IDPs (Vg/Vp) in European ancestry. O indicates significance and X indicates non-significance. B, Estimated heritability of behavior phenotypes in European ancestry. C, SNP heritability point estimates and standard errors (SE) of brain imaging and behavior phenotypes. SNP-based heritability is defined as the proportion of variance in a specific trait or characteristic within a population that can be explained by genetic variations known as single nucleotide polymorphisms (SNPs). In the ABCD youth sample, SNP-based heritability was estimated using a linear mixed model framework, where all common SNPs were fitted simultaneously as random effects to assess the proportion of variance in traits explained by these genetic variations.

Among the structural MRI IDPs, cortical volume was the most heritable (0.27), while curvature was the least (0.22). Among the imaging modalities tested, the volumes of structural MRI IDPs exhibited the highest proportions of heritable brain IDPs. Of the diffusion MRI IDPs, tractography-based IDPs (i.e., dMRI streamline count; 0.19) showed lower heritability than diffusion tensor-based IDPs (i.e., dMRI FA; 0.22), which is consistent with the findings from the adult’s brain in the UK Biobank^8^. While the average heritability of task fMRI IDPs was comparable to that of structural MRI IDPs (with mean heritability estimates being 0.22 for the MID task, 0.24 for the N-back task, and 0.25 for the SST task), the total number of heritable task fMRI IDPs was relatively smaller compared to other brain modalities.

Within the non-brain phenotypes, neurocognition phenotypes showed highest heritability than other psychological measures, followed by behavioral problem measures.

### Association of GPS with Cognitive, Behavioral, and Psychological Outcomes

We tested bivariate correlation between the 14 GPSs and their related phenotypes available in the study dataset of 6,555 European-ancestry and 8,620 multiethnic individuals including covariates (**Table S6**). Among the tested GPSs, the GPSs of cognition-related traits (e.g., cognitive performance, educational attainment, IQ) showed relatively large effect sizes (β) and explained a greater amount of variance of their target outcome compared to other types of outcomes. For instance, IQ GPS explained 18.0% of the variance (adjusted *R^2^*) of the crystalized intelligence scores from NIH Toolbox and showed effect size (β) of 0.286 in European-ancestry children (*P* < 0.0001). Furthermore, BMI GPS explained a relatively high proportion of variation in BMI (β = 0.299; *P* < 0.0001; adjusted *R^2^*= 0.130), followed by the GPSs for psychopathology, such as ADHD, depression, MDD, insomnia, and PTSD with the adjusted *R^2^* ranging from 0.005 to 0.030 (*P* < 0.05). The GPSs of happiness-related traits (i.e., subjective wellbeing, general happiness, happiness-health, happiness-life) showed no significant associations with the variables measuring positive affect of children (i.e., positive emotions and affective well-being in past weeks).

The results in multiethnic children were similar, but the overall variances explained by GPS and covariates were higher than those in European-ancestry children, possibly owing to the increased power. The proportion of variance of cognitive phenotypes explained by the GPSs of cognition-related traits and covariates ranged from 17.0% to 27.4%, and, for BMI, the proportion explained was 15.7% (*P* < 0.0001).

### Multivariate associations among Genome-wide Polygenic Scores (GPSs), Phenotypes, and Brain IDPs

We performed 2-block SGCCA to evaluate the multivariate covariation patterns between the GPSs of multiple complex traits and brain IDPs from 7 neuroimaging modalities. We anticipated identifying significant components of covariation from the CCA results if distributed effects of the GPSs are jointly observed across brain IDPs of a specific modality.

The analyses were primarily performed in the children with European-ancestry, and then were repeated in the entire participants with diverse ancestries for increased statistical power. The analyses identified significant covariation patterns of the GPSs with the brain IDP blocks of structural MRI, diffusion MRI (FA), and N-back task-based functional MRI, indicating significant covariation patterns among these brain IDP-GPS blocks (**Table S7)**. The number of estimated components were empirically determined by maximizing the explained variances of the overall block. Of note, the estimated components of 2-block SGCCA between phenotype and GPS blocks were significant.

To further investigate the covariation patterns of brain IDP-GPS on the expression of phenotypes in preadolescent children, we conducted GPS-brain-phenotype SGCCA (e.g., 3-block CCA) with same sets of GPSs and phenotype blocks, but different brain imaging modalities. Across the 7 sets of 3-block CCA analyses (sMRI, dMRI (streamline count, FA), resting-state fMRI, MID task-based fMRI, SST task-based fMRI, and emotional version of N-back task-based fMRI), all the first two CCA components were significant after FDR correction. It is notable that the GPSs of cognition, such as EA, CP, and IQ showed consistently positive loadings across all the 3-block CCA analyses. The consistent significance of the CCA loadings indicated that the polygenic signals of cognitive traits might have a critical role in the brain and the behaviors. Full results of statistical significance and correlation efficient of each CCA component are available in **Table S7 and Table S8**, respectively.

#### Brain Morphometry (sMRI)

Component-level results: we performed 2-block and 3-block SGCCA on 5,602 European-ancestry individuals’ GPSs, phenotype, and brain morphometric data based on T1-weighted MRI data, including cortical area, volume, thickness, and mean curvature. The independent set of 1,365 individuals with European ancestry was used for hyperparameter tuning of CCA. In 2-block GPS-sMRI CCA, we observed one significant component of covariation (*cov*=1.658; *r*=0.163; *P_fdr_* < 0.001) that accounted for 19.4% and 9.4% of the variance in the sMRI and GPS blocks, respectively **(Table S9)**. Within the sMRI block, 148 out of 389 tested IDPs had non-zero loadings, and 140 of them were significant at *P_fdr_* < 0.05, indicating a significant correlation between brain structural features and genomic components.

The 3-block GPS-sMRI-phenotype CCA identified three components significantly at *Pfdr* < 0.001, which explained 23.9%, 16.9%, and 6.2% of variance of the sMRI, GPS, and phenotype blocks, respectively. The loadings of 3 morphometric IDPs were significantly positive in both 2-block and 3-block CCA, which include total gray volume, left ventral DC, and right accumbens area. The results of GPS-sMRI CCA are available in **Figure 2 and Table S10**.

**Figure 2.**
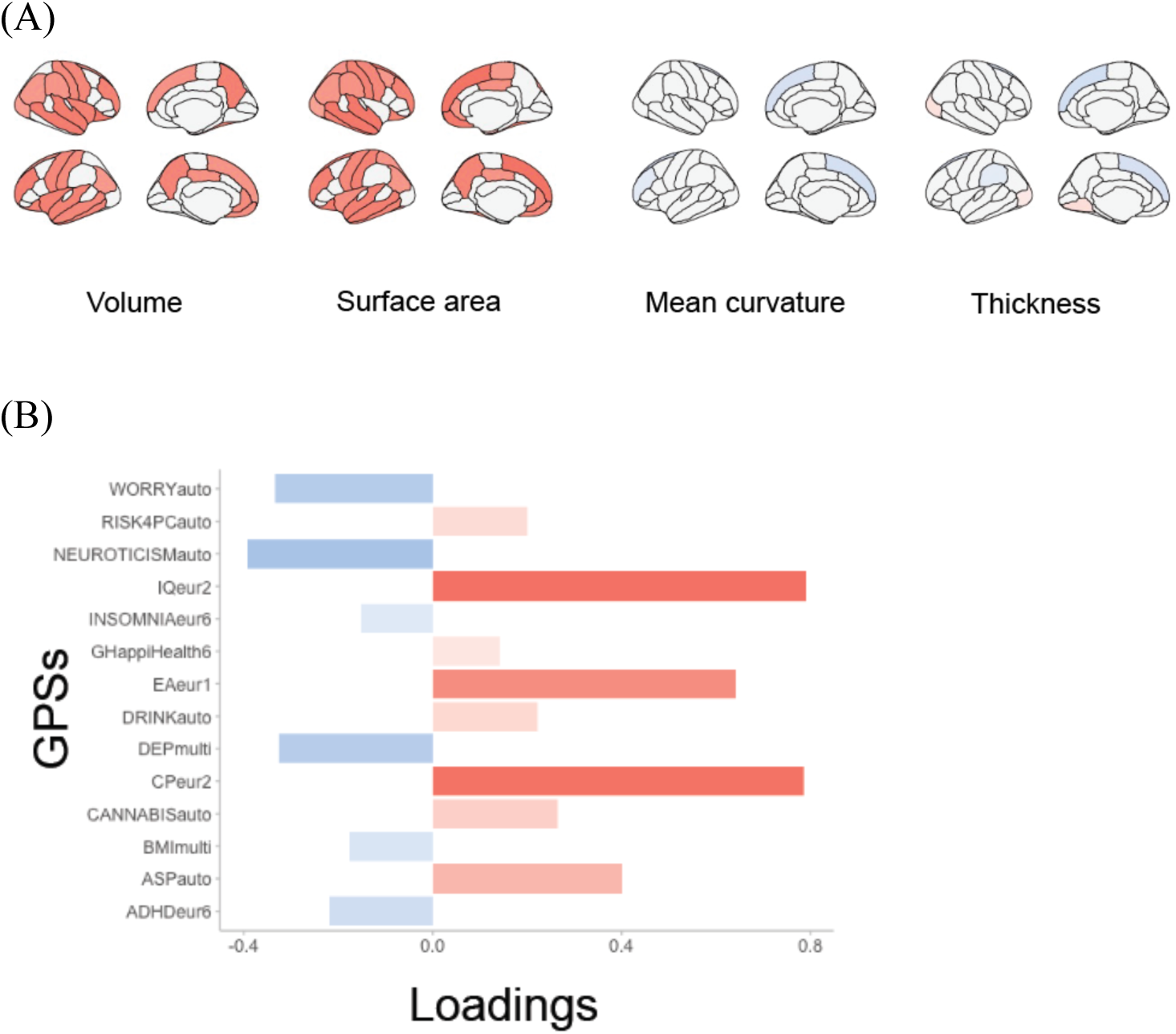
Significant morphometry modes derived from 2 block SGCCA in European ancestry. (A) Significant loadings of the morphometry measures, including regional cortical thickness and cortical surface area (observed for the first significant mode). (B) Significant loadings of GPSs (observed for the first significant mode).

In the GPS block, out of the 29 tested GPSs, the GPSs of 11 traits were identified as significantly associated (*P_fdr_* < 0.05) in the 2-block CCA, and 3 of them (EA, IQ, depression) remained significant in the additional 3-block CCA of European ancestry participants. The GPSs of IQ and EA presented highest positive loadings, whereas the GPS of depression consistently showed significant negative loading.

In the 3-block CCA with sMRI, four phenotypes presented significant covariation with the brain and GPS blocks. Crystallized intelligence showed positive loadings, while mother nervous breakdown problem, CBCL internalizing problem, and father job/fights/police problem presented negative loadings. In the sMRI SGCCA of 7,227 multi-ancestry individuals, the findings from European descents were consistently observed as described in the **Supplementary Results**.

#### Diffusion MRI (dMRI)

##### • Fractional Anisotropy

We conducted 2-block and 3-block SGCCA on 5,842 European-ancestry children’s multi-trait GPSs, phenotype and dMRI data with fractional anisotropy (FA), one of the diffusion tensor imaging metrics, which represents white matter microstructural integrity. The independent validation set of 1,226 individuals of European-ancestry was used for hyperparameter tuning.

In the 2-block GPS-brain CCA of European descents, we identified two significant components of covariation between genomic and dMRI-FA data after conducting permutation testing (*cov_1_* =1.381; *r_1_* =0.102; *P_fdr, 1_* = 0.011; *cov_2_* = -0.942; *r_2_* = -0.173; *P_fdr, 2_* = 0.02). These components cumulatively explained 14.6% and 17.8% of the variance in the dMRI-FA and GPS blocks, respectively. In the dMRI-FA block, we found that 770 out of 2138 tested IDPs had non-zero loadings, and 768 of them were significant at *P_fdr_* < 0.05, indicating a significant correlation between dMRI-FA features and genomic components **(Table S11A).**

The 3-block GPS-brain-phenotype CCA identified three SGCCA components significant at *P_fdr_* < 0.001 after permutation testing, which cumulatively explained 4.20%, 22.43%, 7.32% of the dMRI FA, GPS, and phenotype blocks, respectively, which showed relatively weak proportion of explained variance in brain block compared to other modalities. The loadings of two significant IDPs were consistently found in the 3-block CCA analysis of European descents with positive loadings: connectivity between right hippocampus and right lateral occipital cortex (mean loading = 0.421) and left cerebellum and left amygdala (mean loading = 0.209) (**Table 4**). In the 2^nd^ CCA component, FA measure between left pallidum and paracentral cortex (mean loading = -0.163) was only found significant after FDR correction (*P_fdr_* = 1.38E-02), but it was not found significant in the 3-block CCA of European analysis.

In the GPS block, the GPSs of 3 and 11 traits were identified significantly in the 1^st^ and 2^nd^ components of 2-block CCA analysis, respectively. In the 1^st^ component, the GPSs of BMI (mean loading = 0.650), cannabis use (mean loading = -0.456) and EA (mean loading = -0.568) were significant at *P_fdr_* < 0.05, but none of them were significant in the 3-block CCA of European ancestry. In the 2^nd^ component, the highest positive loadings were observed in the GPSs of neuroticism (mean loading = 0.584), depression (mean loading = 0.560), and schizophrenia (mean loading = 0.507), worrying (mean loading=0.555), cross disorder (mean loading = 0.466), smoking status (mean loading = 0.370), and bipolar disorder (mean loading = 0.312). Interestingly, negative loadings were observed in all the cognition related GPSs (EA, IQ, CP), and subjective wellbeing (mean loading = -0.264). Of them, only the GPSs of depression (mean loading = 0.560) and IQ (mean loading = -0.584) were consistently significant in the 3-block CCA of European descents **(Table S11B).**

The 3-block GPS-brain-phenotype CCA identified five phenotypes significant with the dMRI-FA and GPS blocks. Crystallized intelligence, frequency of physical activity, and UPPS sensation seeking trait presented significant positive loadings, whereas CBCL internalizing problem and Mexican American Cultural Value-family support subscale showed significant negative loadings. We additionally performed 2-block and 3-block dMRI SGCCA of 7,607 multi-ancestry individuals and the results were presented in the **Supplementary Results**.

#### Task-based fMRI

##### • The emotional version of N-back task-based fMRI

We performed 2-block and 3-block SGCCA on 5,549 European-ancestry children’s multi-trait GPSs, phenotypes, and the emotional version of N-back task-based fMRI measuring emotion reactivity and working memory. The independent validation set of 1,141 European-ancestry individuals were used for hyperparameter tuning. The 2-block GPS-brain SGCCA identified one significant component of covariation (*cov*= -0.074; *r*= -0.074; *P_fdr_* < 0.001) that accounted for 13.4% and 10.6% of the variance in the N-back fMRI and GPS blocks, respectively. In the N-back task-based fMRI block, 3 out of 738 tested fMRI IDPs presented non-zero loadings, and only one of them showed a significant correlation (*P_fdr_* < 0.05) with positive loading: mean beta weight of right fusiform area for 0 back condition (mean loading = 0.946) **(Table S12).**

The 3-block GPS-brain-phenotype SGCCA identified three components significant at *P_fdr_* < 0.001 after permutation testing, which cumulatively explained 9.13%, 17.69%, 11.31% of the N-back task-based fMRI, GPS, and phenotype blocks, respectively. The only significant IDP from 2-block CCA was not found significant in 3-block CCAs of European-ancestry participants, but in the additional 2-block and 3-block CCA of the entire multiancestry participants, which results described in the later section.

In terms of polygenic scores, 3 out of the 29 GPSs tested showed non-zero loadings, and only one of them (EA GPS) were significant in the significant 2-block CCA component of European descents. The signal of the EA GPS was robust and consistent, indicating significant covariation patterns with N-back task-based fMRI features with positive loading (mean loading=0.844). The GPS of EA showed consistent and significant covariation (*P_fdr_* < 0.05, mean loading = 0.844) with N-back task-based fMRI features from both 2-block CCA and 3-block CCA. The GPS of depression which presented insignificant non-zero loadings in the 2-block CCA, presented significant negative loadings in the multi-ancestry CCA and 3-block CCA.

The 3-block GPS-brain-phenotype CCA identified four phenotypes significant with the N-back task-based fMRI and GPS blocks. Crystalized intelligence and sensation seeking behavior presented significant positive loadings, whereas mental-health related phenotypes, such as parental externalizing and total problem, showed significant negative loadings.

We additionally performed 2-block and 3-block task-based fMRI SGCCA of 7,137 multi-ancestry individuals and the results were presented in the **Supplementary Results**.

#### Phenotypes

We performed 2-block on 5,549 European-ancestry children’s multi-trait GPSs and behavioral phenotypes (**Figure 3**). All the five components were significant at *P_fdr_* < 0.001 (*cov_1_*= -2.044; *r_1_***=** -0.312; *cov_2_*=0.848; *r_2_***=**0.209) in the European-ancestry individuals, and the 1^st^ component explained the largest amount of covariance, 11.2% and 7.8% in the GPS and phenotype block, respectively.

**Figure 3.**
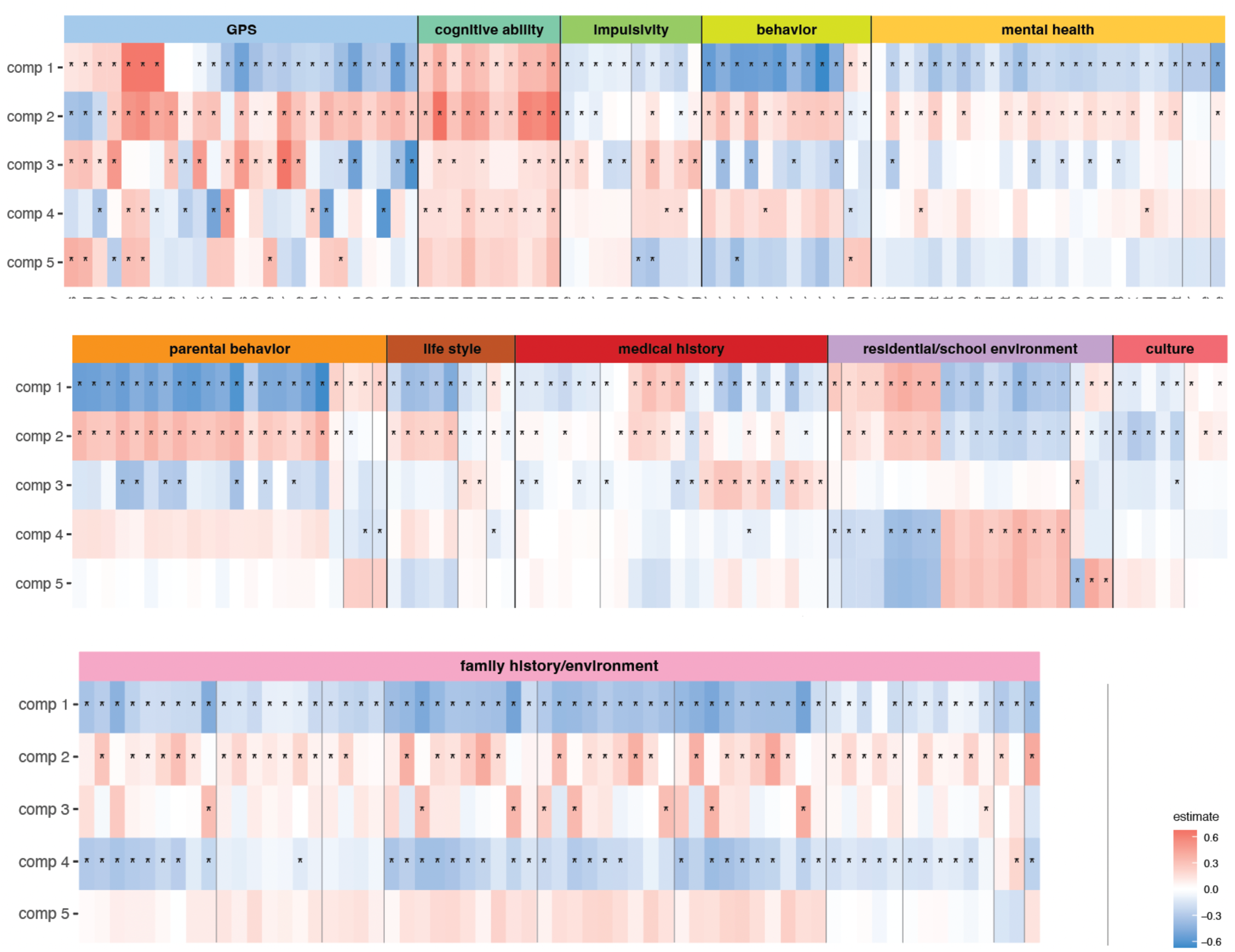
Multivariate associations between phenotypic clusters and genetic predisposition scores in a European ancestry cohort. This figure displays the results of a two-block sparse generalized canonical correlation analysis (SGCCA) illustrating significant associations between phenotype clusters and genetic predisposition scores (GPS). Each row represents a component (comp 1-5) and the columns are phenotypic variables significant in at least one component. Asterisks (*) denote features that are statistically significant within each component, with false discovery rate (FDR)-corrected P-values < 0.05. The color scale reflects the strength and direction of the association, with red indicating a positive estimate and blue a negative estimate.

In the 1^st^ component, 23 out of 25 GPS were significant at *P_fdr_* < 0.05. The GPSs with the highest positive loadings comprised of cognitive traits (IQ, CP, and EA) and happiness traits. The largest negative loadings comprised of several psychiatric outcomes, including depression, smoking status, neuroticism, MDD, and ADHD. The direction of the loadings in the 1^st^ component was consistent with the expected polygenic effects on cognitive ability. In the phenotype block, 195 out of 276 phenotype features were significant at *P_fdr_* < 0.05. The result showed high positive loadings for higher cognitive ability, positive life experiences, family and residential environment, such as NIH Toolbox total composite score, crystallized intelligence, median family income, regional house prices. The largest negative loadings were observed for children’s total behavioral problems, children’s social problems, parental behavioral problems, parental internalizing problems.

In the 2^nd^ component, 24 GPSs and 157 phenotype features were significant at *P_fdr_* < 0.05. In the GPS block, the largest positive loadings were observed for PC of risky behaviors, cannabis use, IQ, CP, and depression and the largest negative loadings were observed for all three tested happiness traits (general happiness, happiness-health, happiness-life). In the phenotype block, the largest positive loadings were observed for parental behavioral problems, parental internalizing problems, parental depressive problems, parental anxious/depressed problems. Of note, largest negative loadings were observed for Mexican American cultural values, positive school environment, family as referent, family support in the phenotype block. The significant results from components 3, 4, and 5 are provided in **Table S13**.

The 3-block CCA analyses revealed that crystallized intelligence, CBCL internalizing problem, UPPS sensation seeking trait, frequency of physical activity, and parental history of job, fights, or police problems are significantly related to both neuroimaging modalities and genetic factors.

The same design of CCA was performed in the 8,619 multiancestry participants, yielding results that were analogous but not identical, as detailed in **Supplementary Results.**

### Predictive performance of the GPS-based models

Based on the significant results of the CCA analysis, we tested predictive power of the PRS for cognitive and psychological outcomes of children. Machine learning trained on GPSs showed better prediction for cognitive outcomes surpassing the non-GPS baseline model (average ΔRMSE_GPS-baseline_ = -0.020; average ΔMAE_GPS-baseline_ = -0.014; average Δ*R^2^*_GPS-baseline_ = 0.035). GPS model for crystallized intelligence score demonstrated superior performance compared to all other models, which gained 5% increase of *R^2^* compared to non-GPS baseline model (RMSE = 0. 863 [SD, 0.019]; MAE = 0.676 [0.015]; *R^2^* = 0.258 [0.020]). GPS models for behavioral problems were marginally better than the baseline (average ΔRMSE = -0.004; average ΔMAE = -0.002; average Δ*R^2^* = 0.008). In contrast, the inclusion of GPSs did not uniformly enhance the performance of models predicting psychiatric disorders (range of Δprecision = -0.010∼0.066; average ΔAUC = - 0.028∼0.020; average Δrecall = -0.133∼0.061). Among the nine classification models, GPS model for ADHD showed the best goodness-of-fit in terms of AUC metrics (AUC = 0.630 [0.019]; ΔAUC = 0.020; **Figure 4B**). The results obtained from multiethnic children were similar to those from European-ancestry children, but with superior performance (**Figure S3**).

**Figure 4.**
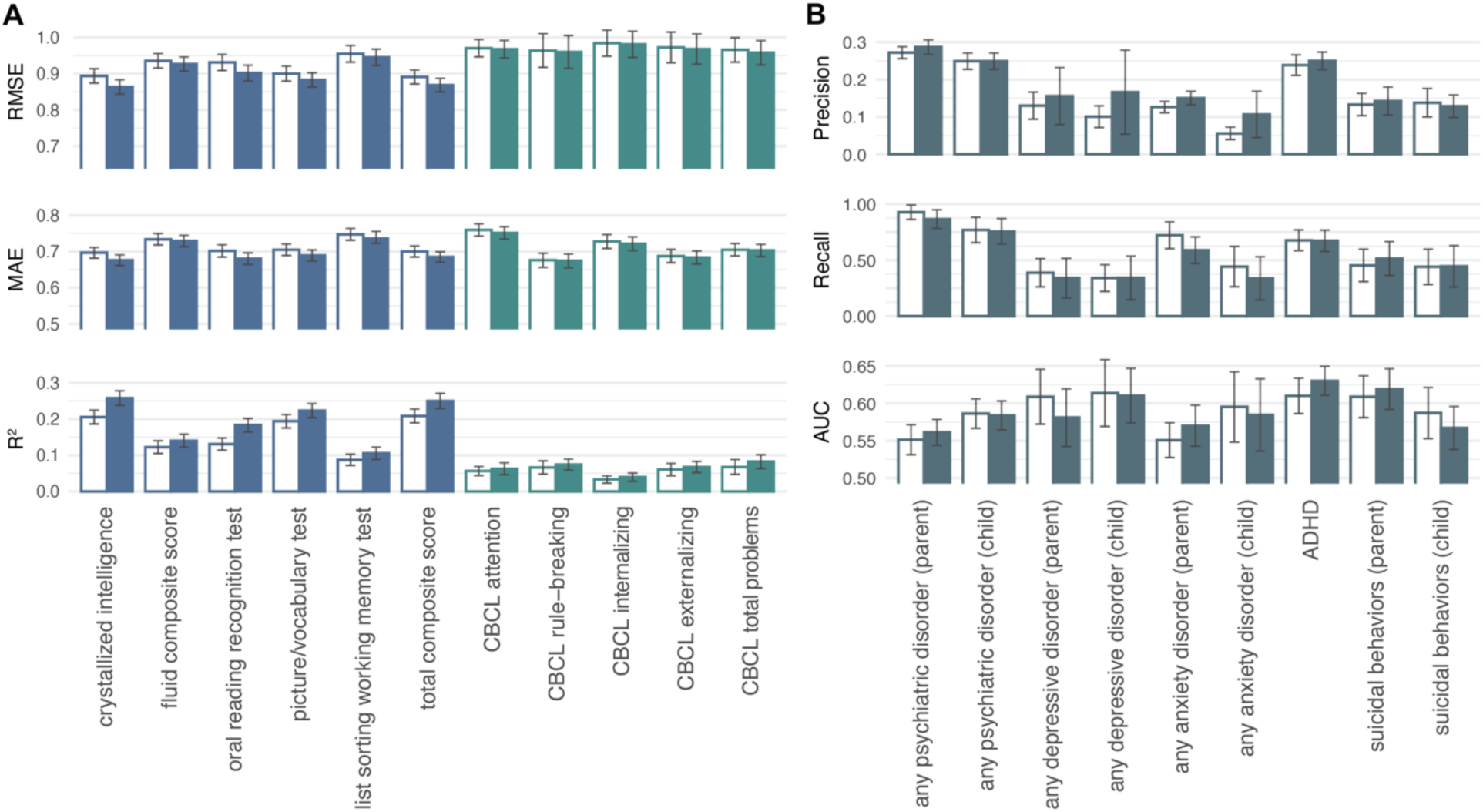
Performance in machine-learning-based association models through GPS integration. Bars without color indicate baseline models, and those filled with color indicate GPS-based models. Error bars show one standard deviation of 5-fold cross-validation metrics. **A**. Root- mean-squared error (RMSE), mean absolute error (MAE), and the variance explained by the predictors (R^2^) of regression models for cognitive and behavioral outcomes. **B**. Precision, recall, and AUC of classification models for psychiatric disorders.

Figure 5 shows the important predictors in the machine learning models. In predicting cognitive outcomes, the models frequently selected GPSs for cognitive performance, IQ, and education attainment as important features. The models for externalizing traits, such as attention problem, rule-breaking behavior, and externalizing behavior, selected the GPSs for ADHD, smoking status, and PC of risky behaviors as significant predictors. For internalizing, the GPSs for neuroticism and depression were important predictors. In the models for psychiatric disorders, the GPSs of cognitive and mental problems, such as ADHD, PTSD, depression, and neuroticism, mainly contributed to classifying the outcome variables. Similar GPSs were shown to be important in multiethnic children (**Figure S4**).

**Figure 5.**
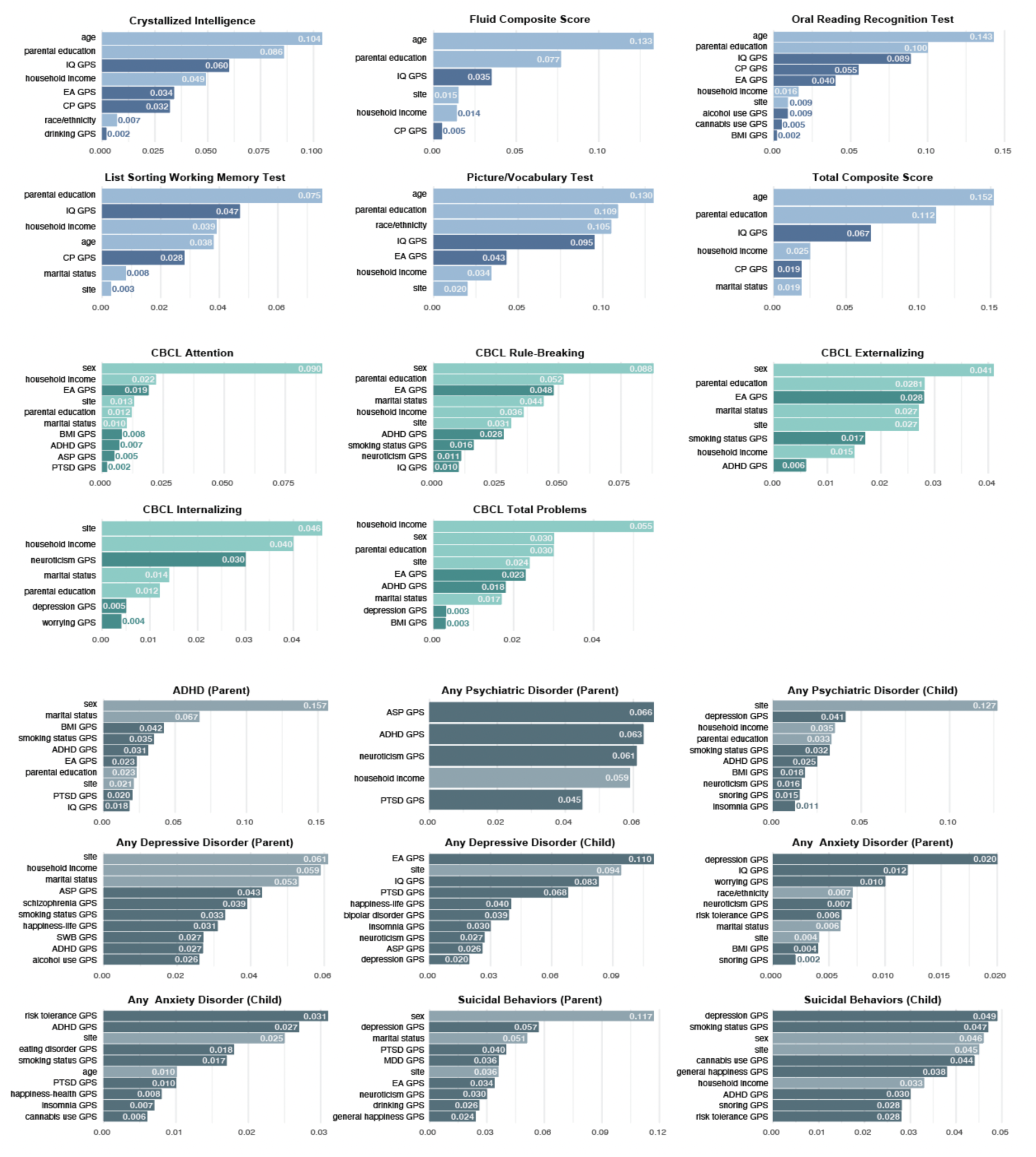
Importance of selected features in GPS-based models in European-ancestry children. Top ten selected predictors are shown with local Shapley values in absolute scales. Features in this figure are restricted to those with absolute values greater than or equal to 0.001. In the bar graph, blue, green, and blue-grey bars respectively represent variables associated with cognitive factors, behavioral problems, and psychiatric disorders or mental issues. The darker shades within each color category signify features derived from GPS data, whereas the lighter shades represent demographic and socioeconomic variables.

## Discussion

In our pursuit to dissect the multivariate relationship of genetics to brain and cognitive development in children, we applied a suite of multivariate analytics including genome-wide polygenic scores (GPS), canonical correlation analyses (CCA), and machine learning approaches. Our main discovery is three-fold. First, SNP-based heritability analyses on brain and behavioral phenotypes in children showed significant genetic contributions, with brain phenotypes exhibiting heightened heritability over behavioral ones. This highlights the genetic foundations steering neurodevelopmental trajectories. Second, Sparse Generalized Canonical Correlation Analysis (SGCCA) with GPSs revealed intricate genetic networks associated with neuroimaging-derived phenotypes (IDPs). This enhances our grasp of the genetic architecture driving cerebral maturation. Notably, GPSs for educational attainment (EA), IQ, and depression were pivotal, with EA and IQ displaying robust positive linkages across diverse brain imaging modalities. Conversely, depression exhibited consistent negative correlations, revealing the genetic intricacies that govern cognitive and mental health attributes. Third, machine learning models trained on the 29 GPSs showed moderate to robust prediction of cognitive outcomes, lower performance for psychopathologies, showing room for improvement towards prognostic utility.

Our heritability analysis identified a number of brain IDPs in preadolescent children that exhibit significant genetic influences across different neuroimaging modalities. Prior heritability analyses in adults showed significant genetic influences on cortical, subcortical regions, and functional connectivity^49-51^. We observed lower heritability of the same brain regions in children compared to adults, aligned with emerging evidence of brain heritability patterns within this developmental stage^9^. This phenomenon indicates that an increase in heritability with age might be a general phenomenon across various traits such as IQ^52^ and could be attributed to multiple factors like decreased environmental variance or gene-environment interactions^6,53^. In contrast to the adult brain, where genetic influences on white matter microstructure surpass those on morphometric features^8^, we also identified relatively stronger genetic influences in morphometric phenotypes, particularly in volumetric measures^54^. This divergence underscores the developmental stage-specific nature of genetic influence and points to a dynamic genetic contribution to neurodevelopmental processes.

Our SGCCA analysis delineated several notable patterns of multivariate covariation among polygenic liabilities of cognitive and mental traits, cognitive and behavioral phenotypes, and brain IDPs of structural MRI, diffusion MRI (FA) and N-back task-based fMRI. Of note, the brain features that emerged as significant in the SGCCA were predominantly those with higher heritability, such as total gray volume (heritability, 0.23), left ventral DC (0.33), and right accumbens (0.30). Cognition-related GPSs showed robust, modality-general covariation patterns with cognitive ability phenotypes, marked by consistently positive loadings. In contrast, GPSs linked to psychiatric conditions and behavioral issues reported by children and parents were associated with negative loadings. These patterns were consistently observed in our 2-block and 3-block SGCCA across the modalities. These findings support the gene-brain pathways underlying the development of cognitive intelligence and psychopathologies.

Particularly within sMRI IDPs, GPS for educational attainment (EA), IQ, and depression significantly covaried with brain volume and surface area, commonly observed in 2-block and 3-block SGCCA. In 2-block CCA, higher GPSs for EA were associated with greater brain volume and surface area, as previously reported^9,55,56^. In stark contrast, GPSs for internalizing behaviors, such as anxiety, neuroticism, and depression, exhibited significant inverse relationships with these brain measures. Of note, a recent CCA study^9^ with the same ABCD data reports no significant associations between brain measures and GPS related to psychopathologies such as bipolar, MDD, OCD, schizophrenia, and ASD. In contrast to this study, our SGCCA approach, incorporating a broader spectrum of GPS, has identified the novel covariation patterns between GPS for psychological traits, including ADHD, depression, and neuroticism, and brain structural features. Our finding highlights the importance of considering a wide spectrum of genetic variables when investigating the neurobiological basis of psychopathology. This approach may lead to enhance our comprehension of the complex relationship between polygenic factors and brain morphology during preadolescent development.

Another novel discovery of our study is the identification of covariation between the depression GPS and brain structural features, such as total grey matter volume. We derived the depression GPS from a large GWAS of broad depression encompassing both clinical diagnosis of MDD and self-reported depressive symptoms^57-59^. In contrast, in our analysis, MDD GPS showed no significant covariation with brain features in contrast to a recent study with an adult sample^60^. While previous studies explored the association between schizophrenia GPS and cortical thickness in from children to adults^16^, both our study and the recent CCA study suggest that this association is not shown in the larger ABCD study sample.

The covariation patterns between white matter microstructure and GPS are novel findings. Within dMRI-FA, both 2-block and 3-block SGCCA pinpointed white matter features of significance including connectivity between the cerebellum and amygdala, hippocampus and lateral occipital gyrus, as well as the paracentral lobule and rostral middle frontal gyrus. This elaborates on previous findings that reported associations between GPSs for educational attainment and IQ with whole-brain FA measures in children^61^.

Within 2-block CCA with dMRI-FA, the second component shows interesting covariation patterns across GPSs. All cognition-related GPSs (e.g., EA, IQ, and CP GPSs) were negatively correlated with the white matter FA measures, whereas the GPSs related to psychiatric disorders (e.g., schizophrenia, bipolar disorder, shared effects on five major psychiatric disorders) and internalizing traits (e.g., neuroticism, depression, worrying) were positively associated. (cf, in the second component, FA measures were not significant at individual feature level, but only at the component level.) These findings suggest the contrasting effects of genetic variants underlying cognitive intelligence and psychopathology on white matter microstructure.

Within task fMRI, significant covariation was found in the right fusiform area for the 0-back condition and EA GPS. This gene-brain activation relationship is particularly compelling, given prior research that underscores the role of fusiform gyrus activation during working memory tasks, especially in relation to dorsolateral prefrontal cortex activation, as a determinant of individual working memory capacity^62^. The SGCCA results suggest a genetic basis for this association, implying that the relationship of the educational attainment GPS with fusiform gyrus activation during working memory tasks may be indicative of broader brain circuitry involvement. It is essential to recognize that the fusiform gyrus’s functional representation is intricately connected to its anatomical pathways extending throughout the brain^63^ (perhaps applicable to other functional domains and circuits^64^). Additionally, the educational attainment GPS reveals a covariation pattern with extensive brain structure and connectivity, hinting that genetic influences on cognitive intelligence might initially impact the structure and connectivity of a broader neural network, which in turn could specifically influence fusiform gyrus activation during working memory tasks. This posited causal pathway, while speculative, calls for rigorous investigation in future studies.

Across different neuroimaging modalities, we observed a consistent set of behavioral phenotypes within our SGCCA analyses. In the first component of the 3-block CCA incorporating dMRI FA, rsfMRI, and the MID task fMRI data, physical activity demonstrated positive covariation with both brain IDPs and GPSs. Conversely, a family history of mental health issues consistently exhibited negative loadings, particularly within the first CCA component for sMRI (e.g., maternal nervous breakdown, paternal employment or legal troubles), dMRI streamline count (e.g., parental employment/legal troubles, alcohol use, depression), and SST fMRI (e.g., parental employment/legal troubles). These bivariate associations between behavior, genetic liability, and the brain have been previously reported^65-69^. Building on this foundation, our findings show the novel tripartite associations among genetics, brain IDPs, and behaviors, suggesting a gene-brain-behavior pathway.

In predicting cognitive outcomes, multivariate patterns of GPSs trained with machine learning showed moderate to good performance. Our full model, incorporating 29 GPSs and socio-economic variables, markedly outperformed our non-GPS baseline model. GPSs contributed to the of crystallized intelligence prediction by *R^2^* of 5.2% and total intelligence prediction by 4.2%. These findings echo our observation of higher heritability of cognitive traits than psychological measures in our analyses and highlight the utility of GPS integration for cognitive ability prediction. Feature importance analysis showed that GPSs for IQ, CP, and EA may be useful in explaining cognitive performance scores. Particularly, EA GPS, which had robust associations with brain IDPs and non-imaging phenotypes in the CCA results, seems to be linked to psychological and behavioral measures as well as neurocognitive measures.

In contrast, in predicting psychological and behavioral outcomes, GPSs made minimal or no contributions. According to the recent studies, prognostic utility of GPS for psychopathologies is doubtful^70-72^. In children, it is perhaps even more so, considering the incomplete development of psychiatric symptoms^73^. Indeed, our prediction results indicated no contributions of GPS to prediction of psychopathologies, except for ADHD where 2% performance gain was found (important GPS included BMI, smoking status, ADHD, and EA GPS). It warrants further investigation to understand the developmental trajectory of GPS-psychopathology associations. Moreover, our predictive analyses suggest that familial (parental mental status, education, family income) and demographic variables (sex) were usually ranked higher than GPSs. This may imply that childhood mental health and cognition are more linked to non-GPS factors such as environmental variables^74-76^.

Despite the varying predictive ability of the GPS model, the primary advantage of the GPS approach is that it provides a simple, interpretable metric that encapsulates an individual’s genetic predisposition for specific traits. This simplicity may enable a useful measure to assess an individual’s genetic liability to complex traits. Nevertheless, it should be noted that GPS typically accounts for a limited portion of genetic liability to psychiatric diseases. Its ability to predict traits is limited when used alone^70^. Our analysis revealed a pronounced disparity between the high SNP-based heritability of brain IDPs and the relatively modest correlations for psychopathology phenotypes using GPS-based predictions. The limited capacity of GPSs at the individual-level traits has been a concern in genetic research. Nevertheless, there is evidence to suggest that, at the population level, GPSs can significantly stratify risk categories, which could be instrumental in targeting interventions more effectively^70,77^.

Despite the limitations, studies have demonstrated that GPSs can facilitate the stratification of populations into distinct risk groups who may benefit more from specific interventions than the general population^78,79^. Furthermore, the predictive strength of GPSs is enhanced when multiple scores are combined with lifestyle or imaging risk factors^80^. Given the intricate interplay and often multiplicative effects of genetic, neurological, and lifestyle factors on complex trait outcomes, GPS information retains its value and importance.

Several limitations of our study warrant consideration. Firstly, we recognize that gene-brain-behavior relationships are likely non-linear, yet our analysis relies on a linear model, which may not fully capture the complexity of these interactions. Secondly, while polygenic scores estimate an individual’s genetic predisposition by summing allele effects across various SNPs, this method may overlook broader gene-brain-behavior patterns that encompass transcriptomic and epigenetic factors. Lastly, our cross-sectional analysis does not allow for causal inferences or the dissection of intricate temporal dynamics among genetic variation, brain phenotypes, and cognitive and behavioral outcomes. Longitudinal studies are needed to shed light on these causal relationships and temporal dynamics, a crucial dimension that our cross-sectional approach does not address.

## Acknowledgement

Data used in the preparation of this article were obtained from the Adolescent Brain Cognitive Development (ABCD) Study (https://abcdstudy.org), held in the NIMH Data Archive (NDA). This is a multisite, longitudinal study designed to recruit more than 10,000 children aged 9–10 and follow them over 10 years into early adulthood. The ABCD Study® is supported by the National Institutes of Health and additional federal partners under award numbers U01DA041048, U01DA050989, U01DA051016, U01DA041022, U01DA051018, U01DA051037, U01DA050987, U01DA041174, U01DA041106, U01DA041117, U01DA041028, U01DA041134, U01DA050988, U01DA051039, U01DA041156, U01DA041025, U01DA041120, U01DA051038, U01DA041148, U01DA041093, U01DA041089, U24DA041123, U24DA041147. A full list of supporters is available at https://abcdstudy.org/federal-partners.html. A listing of participating sites and a complete listing of the study investigators can be found at https://abcdstudy.org/consortium_members/. ABCD consortium investigators designed and implemented the study and/or provided data but did not necessarily participate in the analysis or writing of this report. This manuscript reflects the views of the authors and may not reflect the opinions or views of the NIH or ABCD consortium investigators.

The remaining authors have declared that they have no competing or potential conflicts of interest.

## Data and Code Availability

ABCD https://nda.nih.gov/abcd/request-access

Code Availability: https://github.com/Transconnectome/gps-cca

## Supplementary Materials

### Heritability in multiethnic individuals

Out of the 13,426 IDPs tested, 5,263 IDPs showed significant SNP heritability in 8,620 individuals of multiethnic-ancestry (**Figure S2; Table S2**). SNP heritability in multiethnic participants was low ranging from 0.15 to 0.19. The overall heritability of brain IDPs was higher in multiethnic individuals compared to those of European-ancestry, possibly due to increased sample size. While we found similar patterns of heritability in sMRI and dMRI IDPs between multiethnic and European individuals, we observed that the IDPs of functional connectivity and activation tend to be more heritable in multiethnic individuals (**Figure S2; Table S5**).

### CCA results in multiethnic individuals

#### 1. Structural MRI (sMRI)

In the GPS-sMRI SGCCA of 7,227 multi-ancestry individuals, the findings from European descents were consistently observed. We observed one significant component of covariation (*cov*=1.948; *r*=0.144; *P_fdr_* < 0.001) that accounted for 19.9% and 9.2% of the variance in the sMRI and GPS blocks, respectively. In the sMRI block, all the 389 tested IDPs had non-zero loadings, with 333 of them being significant at *P_fdr_* < 0.05, and 24 were consistently found in the 3-block CCA analysis of the multiancestry participants.

The highest positive loadings in the mutually significant IDPs from the 2-block CCA and 3-block CCA analyses were observed for the following measures: left white matter surface area (mean loading = 0.945), total gray matter volume (mean loading = 0.938), entire cortex volume (mean loading = 0.931), and right (mean loading = 0.930) and left cortex volume (mean loading = 0.929). Conversely, the largest negative loadings were observed for the thickness of right inferior parietal cortex (mean loading = -0.070), right pars triangularis (mean loading = -0.078), mean curve of left inferior temporal cortex (mean loading = -0.087), right lateral orbitofrontal cortex (mean loading = -0.090), left temporal pole (mean loading = -0.104), and the thickness of superior frontal cortex (mean loading = -0.166).

In the GPS block, out of the 29 tested GPSs, 15 traits showed significant loadings at *P_fdr_* < 0.05, with 7 of them consistently being significant in the 3-block CCA analysis of the multiancestry participants, indicating significant covariation with phenotypic features. The GPSs with the highest positive loadings were IQ (mean loading = 0.789), CP (mean loading = 0.785), and EA (mean loading = 0.640). On the other hand, the GPSs with the largest negative loadings were BMI (mean loading = -0.175), ADHD (mean loading = -0.219), depression (mean loading = -0.326), and neuroticism (mean loading = -0.391). Considering substantial genetic overlap between depression, neuroticism, and worrying^29^, our results highlight the high covariation patterns between cognition and those psychiatric internalizing traits with brain morphological features.

#### 2. Diffusion MRI (dMRI) - Fractional Anisotropy

In the 2-block brain-GPS CCA of 7,607 multiethnic participants, we observed a significant component of covariation (*cov*=1.122; *r*=0.100; *P_fdr_* < 0.001) that accounted for 13.86% and 16.3% of the variance in dMRI-FA and GPS blocks, respectively. In the dMRI-FA block, 445 out of 2138 tested IDPs had non-zero loadings, and all of them were significant at *P_fdr_* < 0.05. Among these significant IDPs, 25 were consistently found in the 3-block brain-GPS-phenotype CCA of the multiancestry participants.

Among the mutually significant IDPs from the 2-block CCA and 3-block CCA analyses, the highest positive loadings were observed for the following measures: connectivity between right inferior parietal cortex and right precentral cortex (mean loading = 0.732), left inferior parietal cortex and left putamen (mean loading = 0.730), left inferior parietal cortex and left precentral cortex (mean loading = 0.710), right caudal middle frontal cortex and right inferior parietal cortex (mean loading = 0.673), and left inferior temporal cortex and left precentral cortex (mean loading = 0.664). Conversely, the smallest mean loadings were observed for the connectivity between left thalamus and right cerebellum (mean loading = 0.281), right thalamus and right cerebellum (mean loading = 0.269), right caudate and right cerebellum (mean loading = 0.241), left cerebellum and left amygdala (mean loading = 0.209), and left transverse temporal cortex and left pallidum (mean loading = 0.207), all which loadings are still positive.

In the GPS block, 4 out of 29 tested GPSs were significantly identified in 2-block CCA and all of those 4 GPSs were consistently and significant in the 3-block CCA of multiancestry participants at *P_fdr_* < 0.05. Among significant GPSs, positive mean loading was only observed in the GPS of BMI (mean loading = 0.694), while the GPSs of cannabis use (mean loading = -0.415), CP (mean loading = -0.479), and IQ (mean loading = -0.515) showed negative loadings.

#### 3. Task-based Functional Connectivity (task-based fMRI) - The emotional version of N- back task-based fMRI

The 2-block brain-GPS SGCCA of 7,137 multiancestry children identified one significant component of covariation (*cov*=0.356; *r*=0.081; *P_fdr_* = 0.015) that accounted for 12.9% and 9.9% of the variance in the tfMRI-Nback and GPS blocks, respectively. In the neuroimaging block, 212 out of 882 tested IDPs had non-zero loadings, with 70 of them showing significant correlations (*P_fdr_* < 0.05) with genomic components. The 3-block brain-GPS-phenotype SGCCA identified three components significant at *P_fdr_* < 0.001 after permutation testing, which cumulatively explained 15.77%, 18.31%, 9.63% of the N-back task-based fMRI, GPS, and phenotype blocks, respectively. A total of 7 IDPs were consistently found significant in 2-block and 3-block CCA analysis of the multiancestry participants.

The highest positive loadings in the mutually significant IDPs from the 2-block CCA and 3-block CCA analyses were observed for the following tfMRI-Nback features: mean beta weight of right lingual gyrus for 0 back condition (mean loading = 0.598), left lingual gyrus for 0 back condition (mean loading = 0.593), left superior parietal cortex for 2 back condition (mean loading = 0.588), left fusiform area for 0 back condition (mean loading = 0.575), and right lingual gyrus for emotion condition (mean loading = 0.564). Conversely, the smallest loadings were significantly observed for left (mean loading = 0.110) and right (mean loading = 0.109) caudal middle frontal cortex for 2 back vs. 0 back contrast and left (mean loading = 0.101) and right (mean loading = 0.098) rostral middle frontal cortex for 2 back vs. 0 back contrast, all of which are still positive.

In the GPS block, out of the 29 tested GPSs, the GPSs of 7 traits showed significant associations, with 5 of them consistently being significant in the 3-block CCA of multiancestry participants at *P_fdr_* < 0.05. The GPSs of IQ (mean loading = 0.728), CP (mean loading = 0.729), and EA (mean loading = 0.593) exhibited the highest positive loadings, while the GPSs of depression (mean loading = -0.372), worrying (-0.353), and neuroticism (mean loading = -0.421) significantly showed the largest negative loadings. These findings suggest significant positive associations between the genomic components of cognitive abilities (CP, IQ, and EA) with N-back task-based fMRI features, while genetic influences of internalizing symptoms (e.g., depression, worrying and neuroticism) indicated the opposite covariation patterns.

#### 4. Phenotype

The 2-block phenotype-GPS CCA of 8,619 multiancestry children identified all five components significant at *P_fdr_* < 0.001 (*cov_1_*=1.120; *r_1_***=**0.333; *cov_2_*= -0.395; *r_2_***= -**0.164). The 1^st^ component accounted for 8.7% and 3.2% of the variance in the GPS and phenotype blocks, respectively. In the GPS block of the 1^st^ component, 8 GPSs showed significant non-zero loadings at *P_fdr_* < 0.05. In comparison to the results obtained from the European-ancestry children, the multi-ancestry analysis revealed similar significant phenotypes are associated with GPSs. However, the signs of the loadings for these significant GPSs were found to be opposite in the 1^st^ component. The IQ, CP, and EA GPSs had the largest negative loadings, and smoking status, depression, BMI, and neuroticism GPSs had the largest positive loadings. In the phenotype block, 34 out of 276 tested phenotypes showed significant non-zero loadings at *P_fdr_* < 0.05. The largest positive loadings were observed for NIH Toolbox scores, such as total composite score, crystallized intelligence, and reading ability, and the largest negative loadings were observed for children’s and parent’s problematic behaviors. These variables were shown to be associated also with neuroimaging modalities in 3-block CCA analyses. The strong significance of CCA components was consistently observed in the multiethnic analysis of 8,619 individuals and the results were provided in the **Table S7.**

### Predictive performance of the GPS-based models in multiethnic individuals

Similar results have been observed in European and multiethnic individuals. Machine learning trained on GPSs showed better prediction for cognitive outcomes surpassing the non-GPS baseline model (average ΔRMSE_GPS-baseline_ = -0.012; average ΔMAE_GPS-baseline_ = -0.015; average Δ*R^2^*_GPS-baseline_ = 0.025; **Figure S3**). GPS model for crystallized intelligence score demonstrated superior performance compared to all other models, which gained 4% increase of *R^2^* compared to non-GPS baseline model (RMSE = 0.808 [SD, 0.016]; MAE = 0.631 [0.012]; *R^2^* = 0.348 [0.019]). GPS models for behavioral problems were marginally better than the baseline (average ΔRMSE = - 0.004; average ΔMAE = ∼0; average Δ*R^2^*= 0.008). In contrast, the inclusion of GPSs did not uniformly enhance the performance of models predicting psychiatric disorders (range of Δprecision = -0.003∼0.023; ΔAUC = -0.008∼0.035; Δrecall = -0.133∼0.061). Among the nine classification models, GPS model for ADHD showed the best goodness of fit in terms of AUC metrics (AUC = 0.630 [0.019]), but GPS model for child-reported suicidal behavior gain the most increase of AUC compared to non-GPS baseline model (ΔAUC = 0.035).

Figure S4 shows the important predictors in the machine learning models. In predicting cognitive outcomes, the models frequently selected GPSs for cognitive performance, IQ, and education attainment as important features. The models for externalizing traits, such as attention problem, rule-breaking behavior, and externalizing behavior, selected the GPSs for ADHD, smoking status, and BMI as significant predictors. For internalizing, the GPSs for neuroticism and depression were important predictors. In the models for psychiatric disorders, the GPSs of cognitive and mental problems, such as ADHD, PTSD, depression, and smoking status, mainly contributed to classifying the outcome variables.

## Figures S1 to S4

**Supplementary Figure 1. Estimated heritability of IDPs in multi ancestry.**

**Supplementary Figure 2.**
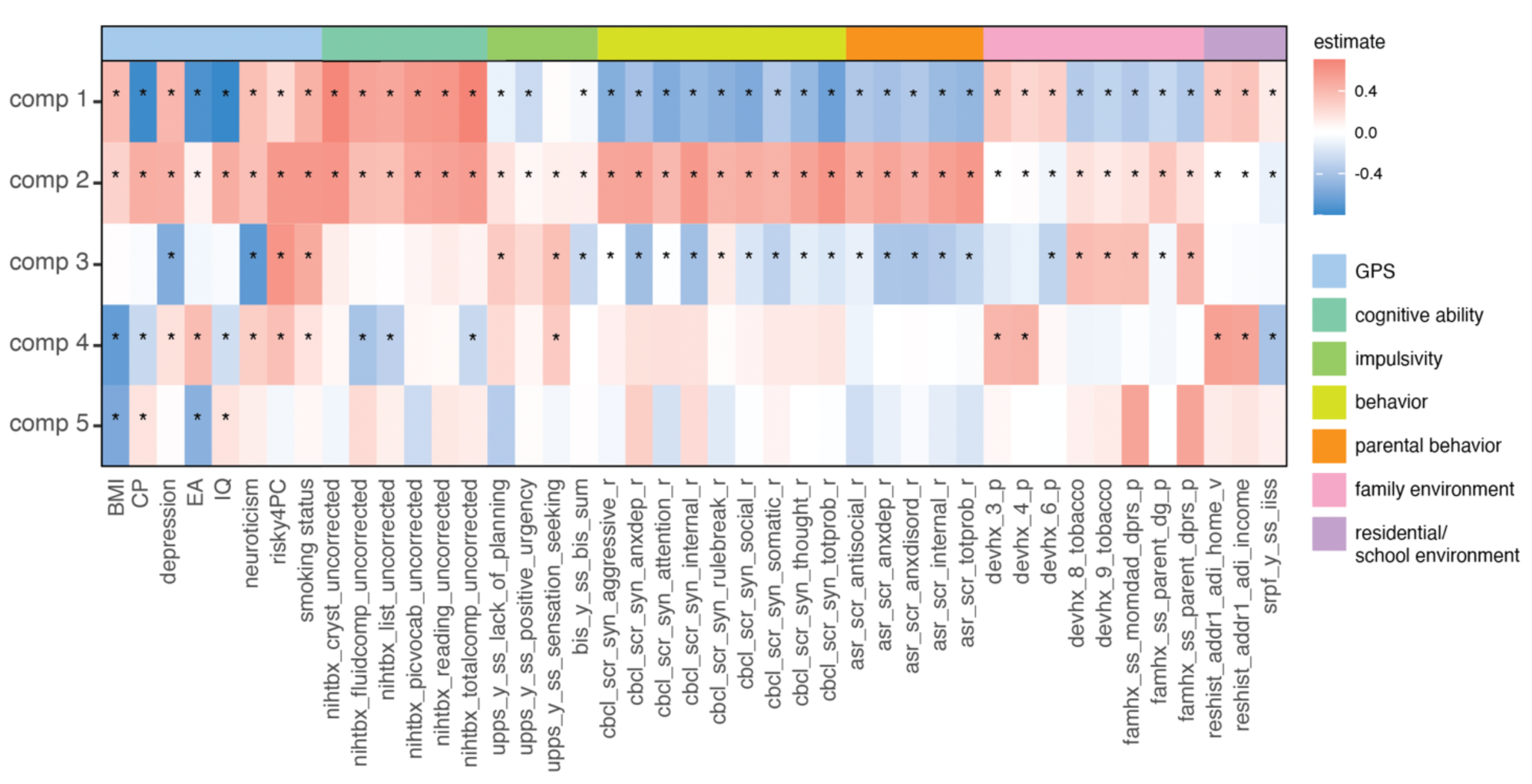
CCA phenotype results in multiethnic individuals.

**Supplementary Figure 3.**
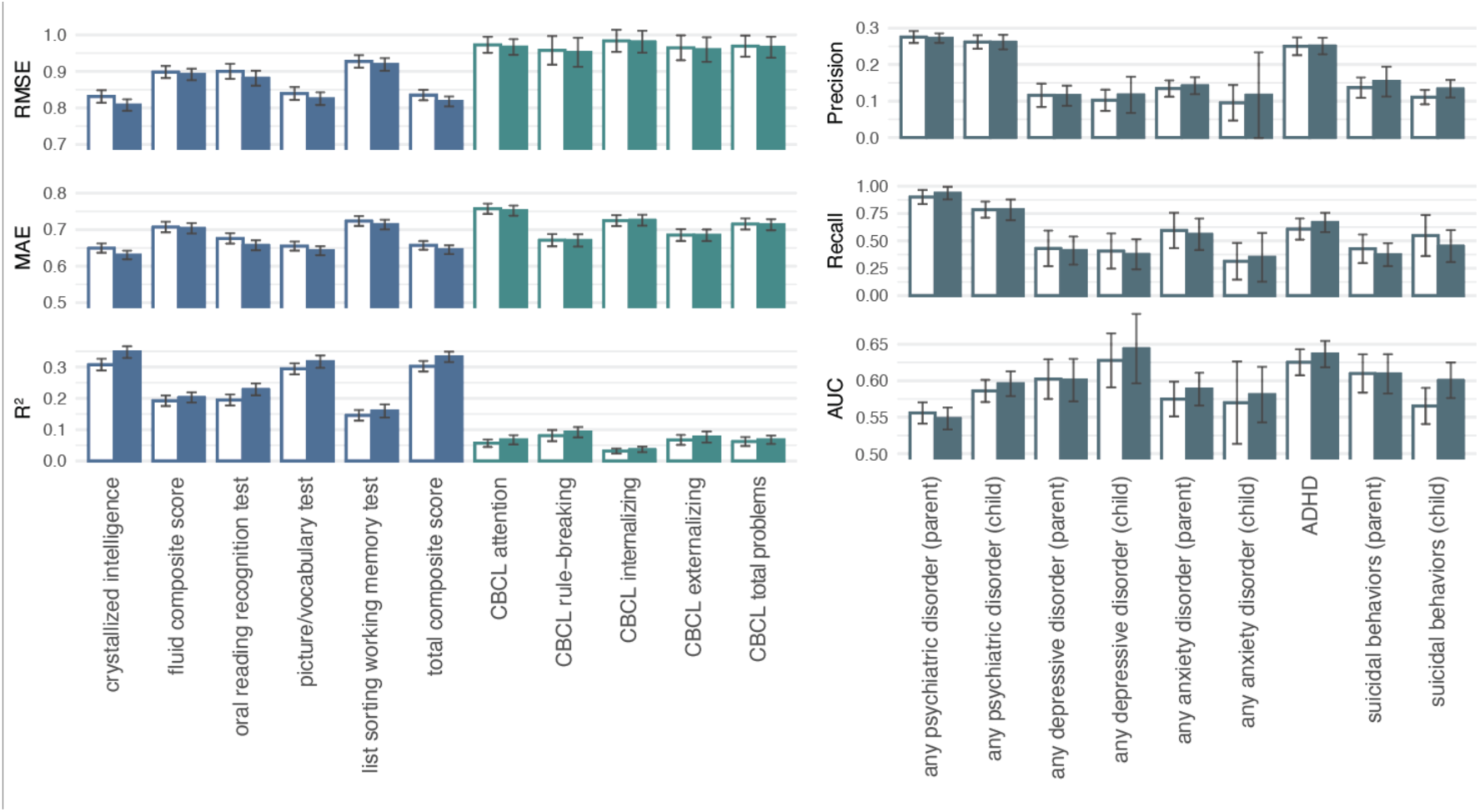
Performance in machine-learning-based association models through GPS integration in multiethnic children.

**Supplementary Figure 4.**
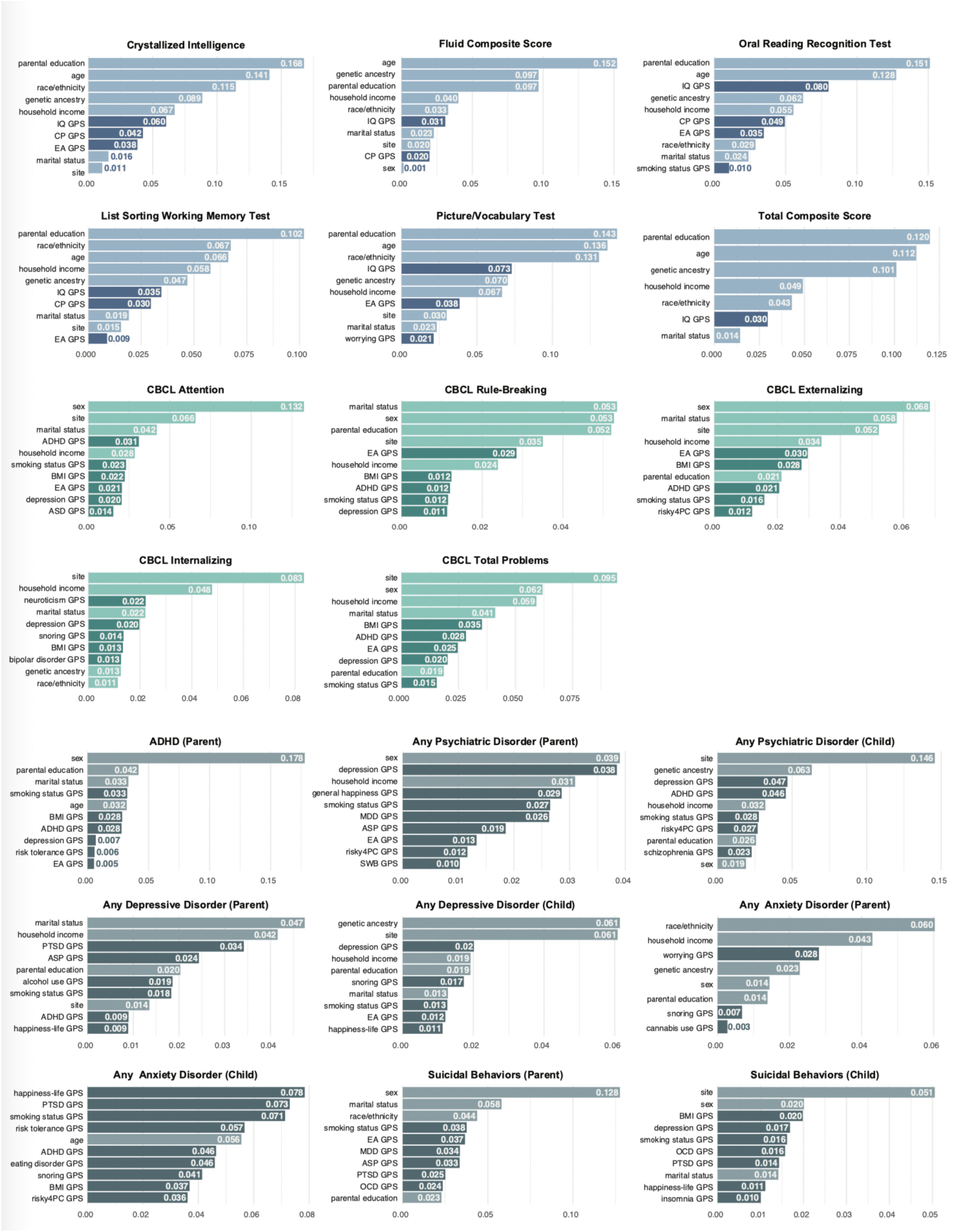
Local Shapley values of selected features in GPS-based models in multiethnic children.

## References

1 Giedd, J. N. et al. Brain development during childhood and adolescence: a longitudinal MRI study. Nature Neuroscience 2, 861–863 (1999). 10.1038/13158

2 Bethlehem, R. A. et al. Brain charts for the human lifespan. Nature 604, 525–533 (2022).

3 Zhao, B. et al. Large-scale GWAS reveals genetic architecture of brain white matter microstructure and genetic overlap with cognitive and mental health traits (n = 17,706). Molecular Psychiatry 26, 3943–3955 (2021). 10.1038/s41380-019-0569-z

4 Maglanoc, L. A. et al. Brain Connectome Mapping of Complex Human Traits and Their Polygenic Architecture Using Machine Learning. Biological Psychiatry 87, 717–726 (2020). 10.1016/j.biopsych.2019.10.011

5 Plomin, R. & Von Stumm, S. The new genetics of intelligence. Nature Reviews Genetics 19, 148–159 (2018). 10.1038/nrg.2017.104

6 Bergen, S. E., Gardner, C. O. & Kendler, K. S. Age-related changes in heritability of behavioral phenotypes over adolescence and young adulthood: a meta-analysis. Twin Research and Human Genetics 10, 423–433 (2007).

7 Brouwer, R. M. et al. Genetic influences on individual differences in longitudinal changes in global and subcortical brain volumes: Results of the ENIGMA plasticity working group. Human brain mapping 38, 4444–4458 (2017).

8 Elliott, L. T. et al. Genome-wide association studies of brain imaging phenotypes in UK Biobank. Nature 562, 210–216 (2018). 10.1038/s41586-018-0571-7

9 Fernandez-Cabello, S. et al. Associations between brain imaging and polygenic scores of mental health and educational attainment in children aged 9–11. NeuroImage 263, 119611 (2022). 10.1016/j.neuroimage.2022.119611

10 Auton, A. et al. A global reference for human genetic variation. Nature 526, 68–74 (2015). 10.1038/nature15393

11 Conomos, M. P., Miller, M. B. & Thornton, T. A. Robust Inference of Population Structure for Ancestry Prediction and Correction of Stratification in the Presence of Relatedness. Genetic Epidemiology 39, 276–293 (2015). 10.1002/gepi.21896

12 Conomos, P., Matthew, Reiner, P., Alexander, Weir, S., Bruce & Thornton, A., Timothy. Model-free Estimation of Recent Genetic Relatedness. The American Journal of Human Genetics 98, 127–148 (2016). 10.1016/j.ajhg.2015.11.022

13 Zheng, X. et al. A high-performance computing toolset for relatedness and principal component analysis of SNP data. Bioinformatics 28, 3326–3328 (2012). 10.1093/bioinformatics/bts606

14 Demontis, D. et al. Discovery of the first genome-wide significant risk loci for attention deficit/hyperactivity disorder. Nature Genetics 51, 63–75 (2019). 10.1038/s41588-018-0269-7

15 Lee, J. J. et al. Gene discovery and polygenic prediction from a genome-wide association study of educational attainment in 1.1 million individuals. Nature Genetics 50, 1112–1121 (2018). 10.1038/s41588-018-0147-3

16 Wray, N. R. et al. Genome-wide association analyses identify 44 risk variants and refine the genetic architecture of major depression. Nature Genetics 50, 668–681 (2018). 10.1038/s41588-018-0090-3

17 Jansen, P. R. et al. Genome-wide analysis of insomnia in 1,331,010 individuals identifies new risk loci and functional pathways. Nature Genetics 51, 394–403 (2019). 10.1038/s41588-018-0333-3

18 Savage, J. E. et al. Genome-wide association meta-analysis in 269,867 individuals identifies new genetic and functional links to intelligence. Nature Genetics 50, 912–919 (2018). 10.1038/s41588-018-0152-6

19 Nievergelt, C. M. et al. International meta-analysis of PTSD genome-wide association studies identifies sex- and ancestry-specific genetic risk loci. Nature Communications 10 (2019). 10.1038/s41467-019-12576-w

20 Howard, D. M. et al. Genome-wide meta-analysis of depression identifies 102 independent variants and highlights the importance of the prefrontal brain regions. Nature Neuroscience 22, 343–352 (2019). 10.1038/s41593-018-0326-7

21 Shen, H. et al. Polygenic prediction and GWAS of depression, PTSD, and suicidal ideation/self-harm in a Peruvian cohort. Neuropsychopharmacology 45, 1595–1602 (2020). 10.1038/s41386-020-0603-5

22 Locke, A. E. et al. Genetic studies of body mass index yield new insights for obesity biology. Nature 518, 197–206 (2015). 10.1038/nature14177

23 Akiyama, M. et al. Genome-wide association study identifies 112 new loci for body mass index in the Japanese population. Nature Genetics 49, 1458–1467 (2017). 10.1038/ng.3951

24 Walters, R. K. et al. Transancestral GWAS of alcohol dependence reveals common genetic underpinnings with psychiatric disorders. Nature Neuroscience 21, 1656–1669 (2018). 10.1038/s41593-018-0275-1

25 Grove, J. et al. Identification of common genetic risk variants for autism spectrum disorder. Nature Genetics 51, 431–444 (2019). 10.1038/s41588-019-0344-8

26 Karlsson Linnér, R., et al. Genome-wide association analyses of risk tolerance and risky behaviors in over 1 million individuals identify hundreds of loci and shared genetic influences. Nature Genetics 51, 245–257 (2019). 10.1038/s41588-018-0309-3

27 Stahl, E. A. et al. Genome-wide association study identifies 30 loci associated with bipolar disorder. Nature Genetics 51, 793–803 (2019). 10.1038/s41588-019-0397-8

28 Pasman, J. A. et al. GWAS of lifetime cannabis use reveals new risk loci, genetic overlap with psychiatric traits, and a causal effect of schizophrenia liability. Nature Neuroscience 21, 1161–1170 (2018). 10.1038/s41593-018-0206-1

29 Consortium, C.-D. G. o. t. P. G. Identification of risk loci with shared effects on five major psychiatric disorders: a genome-wide analysis. The Lancet 381, 1371–1379 (2013). 10.1016/s0140-6736(12)62129-1

30 Watson, H. J. et al. Genome-wide association study identifies eight risk loci and implicates metabo-psychiatric origins for anorexia nervosa. Nature Genetics 51, 1207–1214 (2019). 10.1038/s41588-019-0439-2

31 Nagel, M. et al. Meta-analysis of genome-wide association studies for neuroticism in 449,484 individuals identifies novel genetic loci and pathways. Nature Genetics 50, 920–927 (2018). 10.1038/s41588-018-0151-7

32 Arnold, P. D. et al. Revealing the complex genetic architecture of obsessive–compulsive disorder using meta-analysis. Molecular Psychiatry 23, 1181–1188 (2018). 10.1038/mp.2017.154

33 Ruderfer, D. M. et al. Genomic Dissection of Bipolar Disorder and Schizophrenia, Including 28 Subphenotypes. Cell 173, 1705-1715.e1716 (2018). 10.1016/j.cell.2018.05.046

34 Lam, M. et al. Comparative genetic architectures of schizophrenia in East Asian and European populations. Nature Genetics 51, 1670–1678 (2019). 10.1038/s41588-019-0512-x

35 Okbay, A. et al. Genetic variants associated with subjective well-being, depressive symptoms, and neuroticism identified through genome-wide analyses. Nature Genetics 48, 624–633 (2016). 10.1038/ng.3552

36 Ruan, Y. et al. Improving polygenic prediction in ancestrally diverse populations. Nat Genet 54, 573–580 (2022). 10.1038/s41588-022-01054-7

37 Ge, T., Chen, C.-Y., Ni, Y., Feng, Y.-C. A. & Smoller, J. W. Polygenic prediction via Bayesian regression and continuous shrinkage priors. Nature Communications 10 (2019). 10.1038/s41467-019-09718-5

38 Tournier, J.-D. et al. MRtrix3: A fast, flexible and open software framework for medical image processing and visualisation. NeuroImage 202, 116137 (2019). 10.1016/j.neuroimage.2019.116137

39 Logan, G. D. On the ability to inhibit thought and action: A users’ guide to the stop signal paradigm. (1994).

40 Cohen, A., Conley, M., Dellarco, D. & Casey, B. The impact of emotional cues on short-term and long-term memory during adolescence. Proceedings of the Society for Neuroscience. San Diego, CA. November (2016).

41 Knutson, B., Westdorp, A., Kaiser, E. & Hommer, D. FMRI visualization of brain activity during a monetary incentive delay task. Neuroimage 12, 20–27 (2000).

42 Karcher, N. R. & Barch, D. M. The ABCD study: understanding the development of risk for mental and physical health outcomes. Neuropsychopharmacology 46, 131–142 (2021). 10.1038/s41386-020-0736-6

43 Weintraub, S. et al. The Cognition Battery of the NIH Toolbox for Assessment of Neurological and Behavioral Function: Validation in an Adult Sample. Journal of the International Neuropsychological Society 20, 567–578 (2014). 10.1017/s1355617714000320

44 Weintraub, S. et al. Cognition assessment using the NIH Toolbox. Neurology 80, S54–S64 (2013). 10.1212/wnl.0b013e3182872ded

45 Tenenhaus, A. et al. Variable selection for generalized canonical correlation analysis. Biostatistics 15, 569–583 (2014).

46 Tenenhaus, M., Tenenhaus, A. & Groenen, P. J. F. Regularized Generalized Canonical Correlation Analysis: A Framework for Sequential Multiblock Component Methods. Psychometrika 82, 737–777 (2017). 10.1007/s11336-017-9573-x

47 Hall, P., Gill, N., Kurka, M. & Phan, W. Machine learning interpretability with h2o driverless ai. H2O. ai (2017).

48 Molnar, C. Interpretable machine learning. (Lulu. com, 2020).

49 Baare, W. F. Quantitative Genetic Modeling of Variation in Human Brain Morphology. Cerebral cortex. 11, 816–824 (2001). 10.1093/cercor/11.9.816

50 Roshchupkin, G. V. et al. Heritability of the shape of subcortical brain structures in the general population. Nature Communications 7, 13738 (2016). 10.1038/ncomms13738

51 Shen, K.-K. et al. Investigating brain connectivity heritability in a twin study using diffusion imaging data. NeuroImage 100, 628–641 (2014). 10.1016/j.neuroimage.2014.06.041

52 Bouchard, T. J. The Wilson effect: the increase in heritability of IQ with age. Twin Research and Human Genetics 16, 923–930 (2013).

53 Schmitt, A., Malchow, B., Hasan, A. & Falkai, P. The impact of environmental factors in severe psychiatric disorders. Frontiers in neuroscience 8, 19 (2014).

54 Bethlehem, R. A. I. et al. Brain charts for the human lifespan. Nature 604, 525–533 (2022). 10.1038/s41586-022-04554-y

55 Merz, E. C. et al. Educational attainment polygenic scores, socioeconomic factors, and cortical structure in children and adolescents. Human Brain Mapping 43, 4886–4900 (2022). 10.1002/hbm.26034

56 Judd, N. et al. Cognitive and brain development is independently influenced by socioeconomic status and polygenic scores for educational attainment. Proceedings of the National Academy of Sciences 117, 12411–12418 (2020). 10.1073/pnas.2001228117

57 Neilson, E. et al. Impact of Polygenic Risk for Schizophrenia on Cortical Structure in UK Biobank. Biological Psychiatry 86, 536–544 (2019). 10.1016/j.biopsych.2019.04.013

58 Alnæs, D. et al. Brain Heterogeneity in Schizophrenia and Its Association With Polygenic Risk. JAMA Psychiatry 76, 739 (2019). 10.1001/jamapsychiatry.2019.0257

59 Kirschner, M. et al. Schizophrenia polygenic risk during typical development reflects multiscale cortical organization. Biological Psychiatry Global Open Science (2022).

60 Schmitt, S. et al. Effects of polygenic risk for major mental disorders and cross-disorder on cortical complexity. Psychological Medicine 52, 4127–4138 (2022). 10.1017/s0033291721001082

61. (!!! INVALID CITATION !!! 53,61).

62 Neta, M. & Whalen, P. J. Individual differences in neural activity during a facial expression vs. identity working memory task. Neuroimage 56, 1685–1692 (2011).

63 Saygin, Z. M. et al. Anatomical connectivity patterns predict face selectivity in the fusiform gyrus. Nature neuroscience 15, 321–327 (2012).

64 Cha, J. et al. Circuit-wide structural and functional measures predict ventromedial prefrontal cortex fear generalization: implications for generalized anxiety disorder. Journal of Neuroscience 34, 4043–4053 (2014).

65 Valkenborghs, S. R. et al. The impact of physical activity on brain structure and function in youth: a systematic review. Pediatrics 144 (2019).

66 Zhang, W., Paul, S. E., Winkler, A., Bogdan, R. & Bijsterbosch, J. D. Shared brain and genetic architectures between mental health and physical activity. Translational psychiatry 12, 428 (2022).

67 Tapert, S. F. & Brown, S. A. Substance dependence, family history of alcohol dependence and neuropsychological functioning in adolescence. Addiction 95, 1043–1053 (2000).

68 Posner, J. et al. Increased Default Mode Network Connectivity in Individuals at High Familial Risk for Depression. Neuropsychopharmacology 41, 1759–1767 (2016). 10.1038/npp.2015.342

69 Hao, X. et al. Stability of Cortical Thinning in Persons at Increased Familial Risk for Major Depressive Disorder Across 8 Years. Biological Psychiatry: Cognitive Neuroscience and Neuroimaging 2, 619–625 (2017). 10.1016/j.bpsc.2017.04.009

70 Wray, N. R. et al. From Basic Science to Clinical Application of Polygenic Risk Scores. JAMA Psychiatry 78, 101 (2021). 10.1001/jamapsychiatry.2020.3049

71 Krapohl, E. et al. Multi-polygenic score approach to trait prediction. Molecular Psychiatry 23, 1368–1374 (2018). 10.1038/mp.2017.163

72 Landi, I. et al. Prognostic value of polygenic risk scores for adults with psychosis. Nature Medicine 27, 1576–1581 (2021). 10.1038/s41591-021-01475-7

73 Solmi, M. et al. Age at onset of mental disorders worldwide: large-scale meta-analysis of 192 epidemiological studies. Molecular Psychiatry 27, 281–295 (2022). 10.1038/s41380-021-01161-7

74 Campbell, O. L., Bann, D. & Patalay, P. The gender gap in adolescent mental health: a cross-national investigation of 566,829 adolescents across 73 countries. SSM-population health 13, 100742 (2021).

75 Farah, M. J. et al. Childhood poverty: Specific associations with neurocognitive development. Brain research 1110, 166–174 (2006).

76 Van Droogenbroeck, F., Spruyt, B. & Keppens, G. Gender differences in mental health problems among adolescents and the role of social support: results from the Belgian health interview surveys 2008 and 2013. BMC psychiatry 18, 1–9 (2018).

77 Pain, O., Gillett, A. C., Austin, J. C., Folkersen, L. & Lewis, C. M. A tool for translating polygenic scores onto the absolute scale using summary statistics. European Journal of Human Genetics 30, 339–348 (2022). 10.1038/s41431-021-01028-z

78 Mavaddat, N. et al. Polygenic Risk Scores for Prediction of Breast Cancer and Breast Cancer Subtypes. Am J Hum Genet 104, 21–34 (2019). 10.1016/j.ajhg.2018.11.002

79 Kullo, I. J. et al. Polygenic scores in biomedical research. Nature Reviews Genetics 23, 524–532 (2022). 10.1038/s41576-022-00470-z

80 Dudbridge, F., Pashayan, N. & Yang, J. Predictive accuracy of combined genetic and environmental risk scores. Genetic epidemiology 42, 4–19 (2018).

